# Reliable sensory processing of superficial cortical interneurons is modulated by behavioral state

**DOI:** 10.1101/2024.10.03.616514

**Authors:** Carolyn G. Sweeney, Maryse E. Thomas, Lucas G. Vattino, Kasey E. Smith, Anne E. Takesian

## Abstract

The GABAergic interneurons within cortical layer 1 (L1) integrate sensory and top-down inputs to modulate network activity and induce the plasticity underlying learning. However, little is known about how sensory inputs drive L1 interneuron activity. We used two-photon calcium imaging to measure the sound-evoked responses of two L1 interneuron populations expressing VIP (vasoactive intestinal peptide) or NDNF (neuron-derived neurotrophic factor) in mouse auditory cortex. We find that L1 interneurons respond to both simple and complex sounds with high trial-to-trial variability. However, these interneurons respond reliably to just a narrow range of stimuli, reflecting selectivity to spectrotemporal sound features. This response reliability is modulated by behavioral state and predicted by the activity of neighboring interneurons. Our data reveal that L1 interneurons exhibit sensory tuning and identify the modulation of response reliability as a potential mechanism by which L1 relays state-dependent cues to shape sensory representations.

## Introduction

Sensory perception is dynamically modulated by behavioral state and learned associations^1,2,3,4^. This modulation is observed in sensory cortex, where an elaborate network of neuromodulatory and top-down projections powerfully filter and modify sensory signals relayed from the thalamus. This network influences moment-to-moment cortical activity and drives long-lasting changes in synaptic connections that underlie behavioral learning^5,6,7,8,9,10,11,12,13,14,15^. How these diverse inputs converge onto specific neuronal types within cortex to mediate perception and learning is still unknown.

Recent work has identified cortical layer 1 (L1) as a junction for bottom-up sensory inputs and top-down modulation^16,17^. Axons arising from thalamic nuclei, neuromodulatory centers, motor, and limbic regions create a dense latticework that characterizes this superficial layer^9,13,14,15,16,17,18,19,20,21,22,23,24,25,26,27,28,29,30,31,32^. L1 is exclusively populated by GABAergic inhibitory interneurons that integrate these diverse inputs^13,15,33^. The two most abundant interneuron subtypes in L1 are characterized by the molecular markers vasoactive intestinal peptide (VIP) and neuron-derived neurotrophic factor (NDNF)^14,34^. These interneurons are known to relay information about behavioral state and outcome. For example, *in vivo* studies have demonstrated that arousal, reward, and punishment activate VIP and NDNF interneurons^10,12,14,21,35,36,37,38^. It is thought that VIP and NDNF interneurons integrate these behavioral signals with sensory information to powerfully modulate cortical activity and plasticity during sensory processing and learning^21,35,36,37,38,39^. However, little is known about how these interneurons are recruited by specific sensory stimuli.

Anatomical studies show that L1 receives robust sensory inputs from both primary and higher-order thalamic nuclei^8,15,33,40,41^. In auditory cortex (ACtx), thalamic projections from the primary auditory thalamus (the ventral division of the medial geniculate body; MGBv) extend to L1 to relay fast, frequency-tuned sound information^15,33,42^. The thin band of MGBv axons in L1 forms a coarse spatial map of sound-frequency representation that mirrors the tonotopy in layers 3/4^15,43^. The importance of these direct thalamocortical projections to L1 has been largely ignored. However, recent results have demonstrated that these MGBv axons within L1 of ACtx form robust monosynaptic connections onto L1 VIP and NDNF interneurons that are comparable in strength to those onto pyramidal neurons^15,33,44^. This connectivity predicts that these L1 interneurons display robust responses to sound and tuning to sound frequency. Indeed, a few studies have observed sound-evoked responses in superficial interneurons^14,21,38,45^. However, the selectivity of the L1 interneuron subtypes to simple and complex sounds in awake animals is not well characterized. Understanding how VIP and NDNF interneurons process sensory inputs during distinct behavioral states will be critical to understanding the function of superficial cortex in state-dependent sensory processing and learning.

In the present study, we used two-photon calcium imaging in the ACtx of awake mice to characterize the responses of L1 VIP and NDNF interneurons to a battery of simple and complex sound stimuli and compare these to L2/3 pyramidal neuron (PYR) responses. We found that VIP and NDNF interneurons responded robustly to sound but showed significantly greater trial-to-trial variability than PYR neurons. Moreover, subsets of VIP and NDNF interneurons showed remarkably narrow tuning to specific simple or complex sound stimuli comparable to PYR neurons. Our results suggest that the sensory processing of these interneurons is shaped by interactions between neighboring L1 interneurons and by distinct behavioral states, such as locomotion. Together, these data reveal that VIP and NDNF interneurons within cortical L1 networks encode the sensory environment in a state-dependent manner.

## Results

### VIP and NDNF interneurons respond to sound but with less trial-to-trial reliability than pyramidal neurons

To compare sound-evoked responses across PYR, VIP, and NDNF neurons, we performed two-photon calcium imaging in the superficial ACtx of mice selectively expressing GCaMP6f in these populations (Fig. 1a-c, Supplementary Fig. 1a). For all experiments, head-fixed mice listened passively to simple and complex auditory stimuli while running at will on a wheel designed to minimize movement-generated sound. We focused our imaging on VIP and NDNF interneurons within L1 and along the L1/2 border, and PYR neurons within L2/3 (see Methods). Neurons within all three populations exhibited sound-evoked fluctuations in calcium activity with various temporal profiles. Subsets of neurons showed short-latency positive responses, delayed prolonged positive responses, or suppression. To capture this response heterogeneity, we characterized each neuron as having an *activated*, *prolonged*, or *suppressed* average response profile to each stimulus type (Supplementary Fig. 1b; Methods).

**Figure 1.**
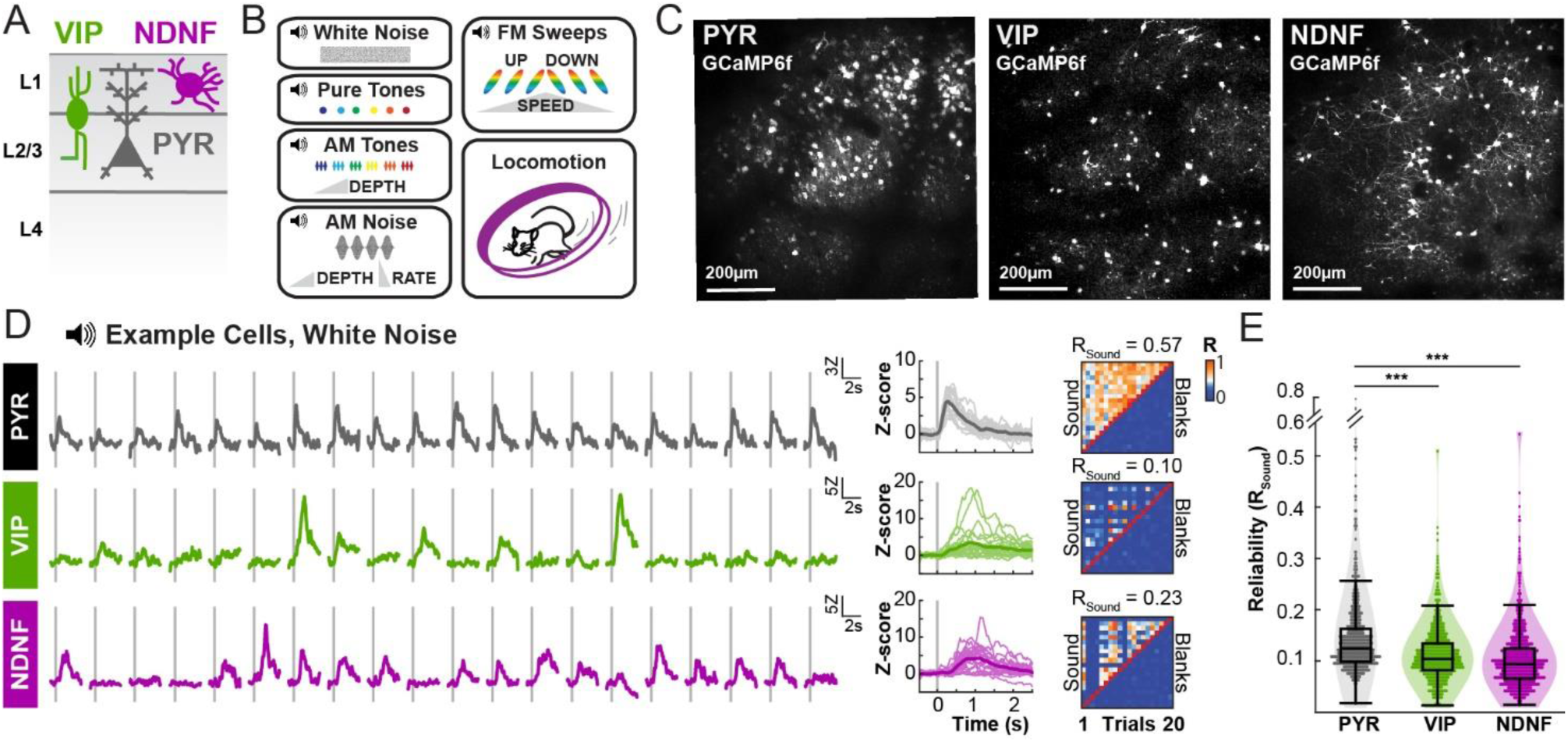
VIP and NDNF interneurons respond to sound but with less trial-to-trial reliability than pyramidal neurons. **(A)** Schematic showing relative position of PYR, VIP, and NDNF populations in the cortex. **(B)** Simple and complex sound stimuli were passively presented to head-fixed mice during two-photon calcium imaging of GCaMP6f in the left auditory cortex. **(C)** Example fields-of-view from each neuronal population. Maximum projection images are shown to highlight differences in density between each population. **(D)** Example single-cell responses to white noise bursts (20 trials) highlighting trial-to-trial response variability. Left, responses to twenty trials of identical white noise burst stimuli. Middle, overlaid single-trial responses with average in bold. Right, Heat map of Pearson’s R calculated for each pair of trials (upper diagonal: sound trials, lower diagonal: blank trials). The average Pearson’s R across all sound trials is the reliability score (R_Sound_) for that neuron. **(E)** Average reliability score (R_Sound_) for each neuronal group. Single points denote mean reliability across auditory stimuli for a single neuron. Mixed-effects model one-way ANOVA with Tukey’s post-hoc test (F_2,2504_ = 77.48, p<0.001), ***p<0.001, n_PYR_ = 694 neurons, n_VIP_ = 773 neurons, n_NDNF_ = 829 neurons.

Subsets of VIP and NDNF interneurons exhibited robust sound-evoked responses to both simple and complex stimuli. However, we were surprised to find that these interneurons displayed high trial-to-trial variability as compared to PYR neurons. Looking at sound-evoked responses across trials during which identical sound stimuli were presented, we observed large stimulus-locked responses during some trials but no responses during other trials (Fig. 1d). We quantified the trial-to-trial reliability of sound-evoked responses by computing the average pairwise Pearson’s correlation, R_Sound_, between responses across all trials with matched sound stimuli (Supplementary Fig. 1c). To detect reliable responses above noise, R_Sound_ was compared to a control distribution of R values generated from blank (silent) trials interleaved with stimulus presentations (^R̅^_Blank_). Only cells with significant R_Sound_ above ^R̅^_Blank_ were considered responsive for further analysis (Fig. 1d).

VIP and NDNF interneurons showed significantly smaller R_Sound_ (Fig. 1e), regardless of auditory stimulus type or response profile (Supplementary Fig. 1d,e). The lower R_Sound_ in the interneuron populations was associated with greater variability in sound response magnitude, but not response width or onset jitter as compared to PYR neurons (Supplementary Fig. 1f-i). Together, these data demonstrate that VIP and NDNF interneurons in ACtx are sound responsive but demonstrate high trial-to-trial response variability.

### VIP and NDNF interneurons show narrow ‘reliable’ response frequency tuning

Previous work shows that L1 of ACtx receives sound frequency-tuned inputs from the primary auditory thalamus^15,43^, suggesting that the interneurons within this layer may display selectivity to sounds of specific frequencies. We compared tonal receptive fields measured from L1 VIP and NDNF interneurons to those of L2/3 PYR neurons, which are known to exhibit frequency tuning^46,47,48,49,50,51^. Within all three populations, we found neurons that responded to pure tone stimuli (32.7% PYR, 25.0% VIP, 17.8% NDNF) with *activated*, *prolonged*, or *suppressed* response types (Fig. 2a). We focused on neurons with *activated* response profiles to assess tuning properties. Subsets of VIP and NDNF interneurons showed selectivity for specific frequencies, with receptive fields that resembled classically tuned PYR neurons (Fig. 2a). We computed the best frequency (BF) for each neuron as the frequency that elicited the strongest response at any sound intensity. Neurons across all populations showed selectivity to a wide range of frequencies. However, the average BF-centered receptive fields of VIP and NDNF interneurons appeared more disordered (Fig. 2b, left) and less sharply tuned to frequency (Fig. 2b, right) as compared to PYR neurons. Consistent with the disordered receptive fields, we measured reduced d′ in VIP and NDNF interneurons, indicating lower receptive field quality^47,52^ (Fig. 2c). However, not all frequency-intensity tone combinations elicited reliable responses in our recordings, and we asked how differences in trial-to-trial reliability across these neuronal populations impacted our measures of receptive field tuning. R_Sound_ was greatest at BF for all three populations (Fig. 2d). For VIP and NDNF interneurons, however, R_Sound_ decreased precipitously to sound frequencies flanking the BF, resulting in just a narrow set of sound stimuli for which these interneurons responded reliably (Fig. 2d). Indeed, when we applied a mask to restrict our receptive fields to only reliable responses, we found that VIP and NDNF interneurons exhibited more narrow tuning curves than PYR neurons (Fig. 2e). Therefore, these interneurons show selectivity for a narrow range of preferred sound stimuli that elicit reliable responses.

**Figure 2.**
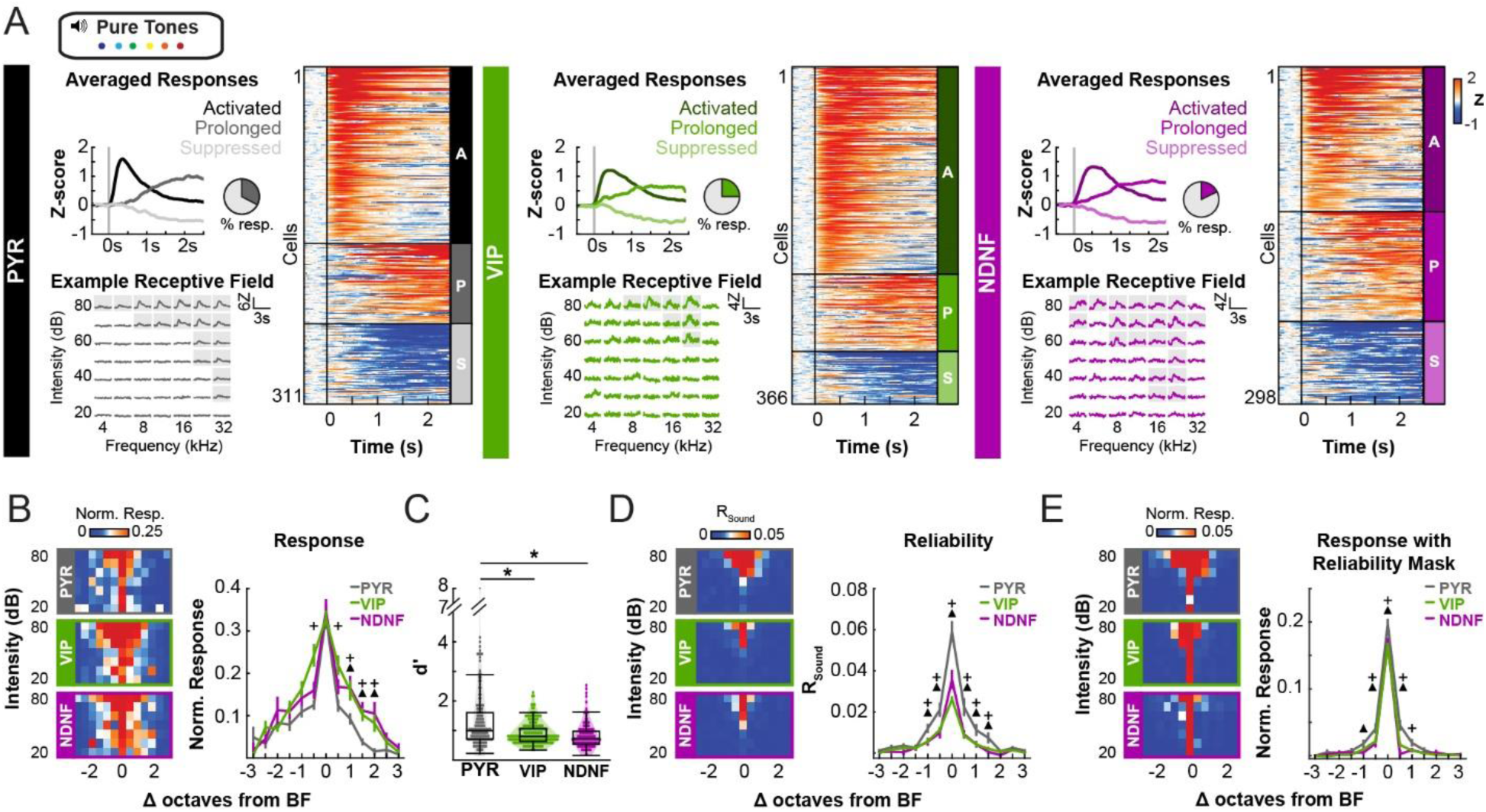
Reliable tonal receptive fields of VIP and NDNF interneurons are narrower than those of PYR. **(A)** For PYR, VIP, and NDNF neuronal populations: top left, average response for all *activated*, *prolonged*, and *suppressed* neurons to pure tones (4-32kHz, 20-80dB). Vertical line denotes sound onset. Pie chart represents the percentage of total neurons responsive to pure tone stimuli. Bottom left, example pure tone receptive field for a single neuron with reliable responses denoted by gray boxes. Right, raster plot showing the mean, Z-scored reliable responses to pure tone stimuli for all responsive neurons. Each row represents a single neuron. Data are sorted by maximal response within *activated* (A), *prolonged* (P), and *suppressed* (S) profiles (n_PYR_ = 311, n_VIP_ = 366, n_NDNF_ = 298). **(B)** Left, average tonal receptive fields for *activated* neurons, normalized to BF. Tonal receptive field contains all sound-evoked responses for each neuron. Right, mean response across all intensities for normalized, *activated* receptive fields. Mixed-effects two-way ANOVA, neuron group x octave bin with post-hoc t-test with Bonferroni correction within octave bin compared to PYR (F_24,6621_ = 2.2046, p=0.0006), ^+^VIP or ^Δ^NDNF significantly different from PYR (F_2,455.2_ ≤ 3.4816, p<0.036), p<0.05, n_PYR_ = 163 neurons; n_VIP_ = 225 neurons; n_NDNF_ = 128 neurons. **(C)** Tonal receptive field organization as measured by d′. Single dots represent d′ from individual *activated* neurons. Mixed-effects one-way ANOVA with Tukey’s post-hoc test (F_2,38.2_ = 14.63, p <0.001), *p<0.05, n_PYR_ = 163 neurons; n_VIP_ = 225 neurons; n_NDNF_ = 128 neurons. **(D)** Average response reliability score (R_Sound_) normalized as in (**B**). Mixed-effects two-way ANOVA, neuron group x octave bin with post-hoc t-test with Bonferroni correction within octave bin compared to PYR (F_24,5866_ = 11.0437, p<0.0001) ^+^VIP or ^Δ^NDNF significantly different from PYR (F_2,651.5_ ≤5.3413, p<0.005), p<0.014, n_PYR_ = 163 neurons; n_VIP_ = 225 neurons; n_NDNF_ = 128 neurons. **(E)** Average tonal receptive field, generated from only reliable responses, normalized to BF. Mixed-effects two-way ANOVA, neuron group x octave bin with post-hoc t-test with Bonferroni correction within octave bin compared to PYR (F_24,6633_ = 2.7155, p<0.0001), ^+^VIP or ^Δ^NDNF significantly different from PYR (F_2,238_ ≤ 3.9766, p <0.02), p<0.026, n_PYR_ = 163 neurons; n_VIP_ = 225 neurons; n_NDNF_ = 128 neurons.

### VIP and NDNF interneurons respond robustly to complex auditory stimuli with diverse response properties

Naturalistic sounds, such as communication calls and environmental sounds, exhibit temporal modulations, including sinusoidal amplitude modulation (SAM) and frequency modulation (FM). We therefore asked how VIP and NDNF interneurons respond to SAM and FM complex sounds as compared to PYR neurons. Subsets of all three populations responded to SAM tones that varied in carrier frequency and modulation depth (40.2% PYR, 25.4% VIP, 23.8% NDNF, Fig. 3a) and SAM broadband white noise that varied in modulation rate and modulation depth (33.4% PYR, 18.7% VIP, 20.2% NDNF, Fig. 3f). Notably, we found that some VIP and NDNF interneurons were well tuned to specific complex sounds (Fig. 3a,f). However, similar to the tonal receptive fields, VIP and NDNF interneurons exhibited more disordered receptive fields for these SAM sound stimuli (Fig. 3b,g, left), broader tuning for SAM carrier frequency and rate (Fig. 3b,g, right), and decreased tuning quality (d′, Fig. 3c,h). Examining response reliability (Fig. 3d,i) again revealed narrower tuning curves for reliable responses recorded in VIP and NDNF interneurons as compared to PYR neurons (Fig. 3e,j).

**Figure 3.**
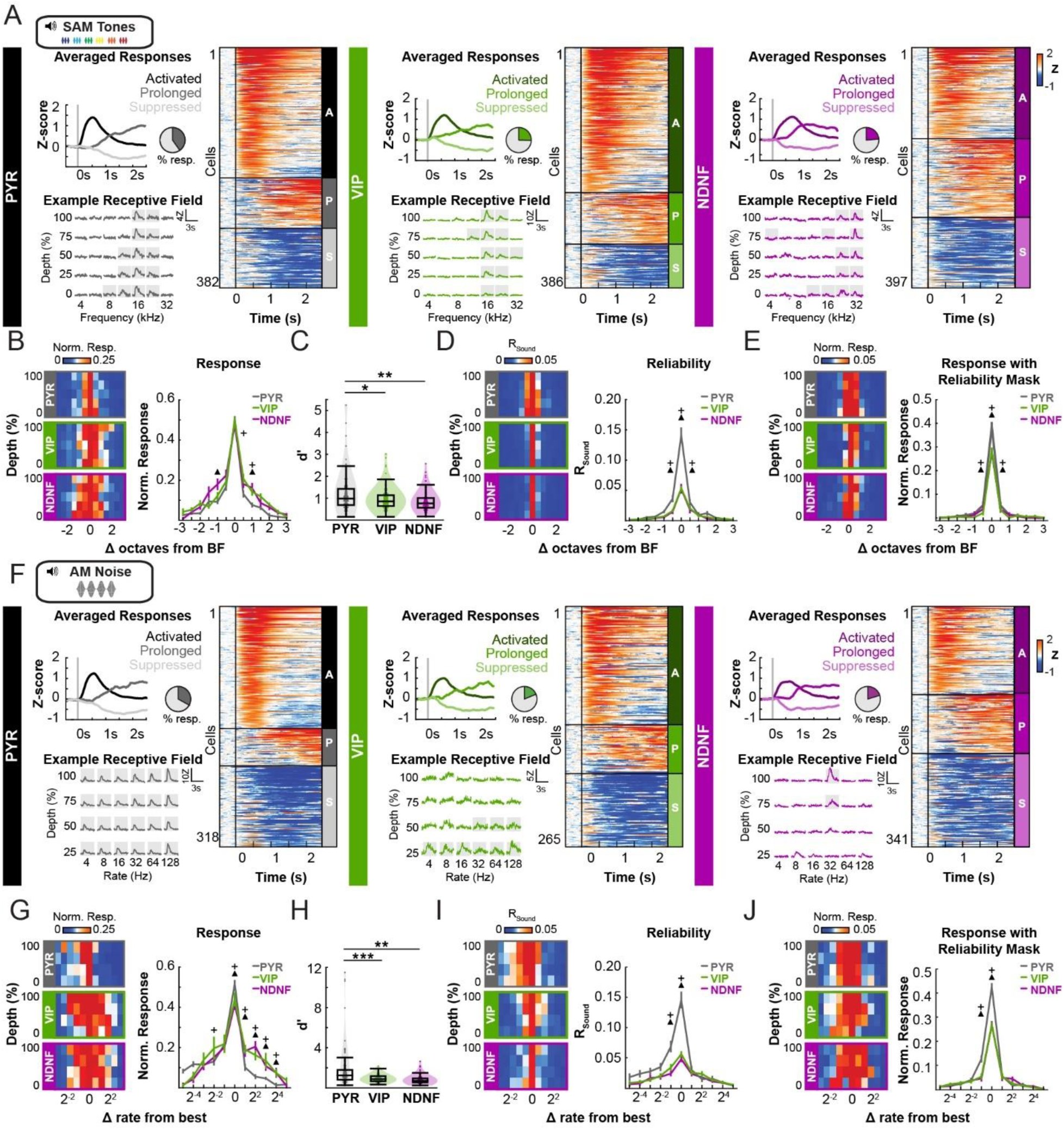
VIP and NDNF neurons respond reliably to a narrow range of complex auditory stimuli. **(A)** For PYR, VIP, and NDNF neuronal populations: top left, averaged responses for *activated*, *prolonged*, and *suppressed* neurons to amplitude modulated tones (SAM tones, 4-32 kHz,10Hz modulation rate, 0-100% modulation depth, 65dB). Vertical line denotes sound onset. Pie chart represents the percentage of total neurons responsive to SAM tones. Bottom left, example SAM tone receptive field for a single neuron with reliable responses denoted by gray boxes. Right, raster plot showing all mean Z-scored reliable responses to SAM tone stimuli. Each row represents a single neuron. Data are sorted by maximal response within *activated* (A), *prolonged* (P), and *suppressed* (S) profiles. (n_PYR_ = 382, n_VIP_ = 386, n_NDNF_ = 397) **(B)** Left, average SAM tone receptive fields for *activated* neurons, normalized to BF. Right, mean response across intensities for normalized, *activated*, receptive fields. Mixed-effects two-way ANOVA, neuron group x octave bin with post-hoc t-test with Bonferroni correction within octave bin compared to PYR (F_24,7641_ = 2.1724, p=0.0008), ^+^VIP or ^Δ^NDNF significantly different from PYR (F_2,283.3_ = 3.950 minimum for significant comparisons, p < 0.02), p<0.04, n_PYR_ = 209 neurons; n_VIP_ = 232 neurons; n_NDNF_ = 153 neurons. **(C)** Receptive field organization as measured by d′. Single dots represent d′ from individual *activated* neurons. Mixed-effects one-way ANOVA with Tukey’s post hoc test (F_2,27.97_ = 9.7756, p =0.014), **p=0.001, *p=0.012, n_PYR_ = 209 neurons; n_VIP_ = 232 neurons; n_NDNF_ = 153 neurons. **(D)** Average response reliability score (R_Sound_) normalized as in (**B**). Mixed-effects two-way ANOVA, neuron group x octave bin with post-hoc t-test with Bonferroni correction within octave bin compared to PYR (F_24,7652_ = 18.3157, p<0.001), ^+^VIP or ^Δ^NDNF significantly different from PYR (F_2,428_ = 4.0159 minimum for significant comparisons, p <0.019), p<0.039, n_PYR_ = 163 neurons; n_VIP_ = 225 neurons; n_NDNF_ = 128 neurons. (**E)** Average receptive field, generated from only reliable responses, normalized as in (**B**). Mixed-effects two-way ANOVA, neuron group x octave bin with post-hoc t-test with Bonferroni correction within octave bin compared to PYR (F_24,6621_ = 2.2046, p=0.0006), ^+^VIP or ^Δ^NDNF significantly different from PYR (F_24,7651_ = 10.1802, p<0.001), ^+^VIP or ^Δ^NDNF significantly different from PYR (F_(2,340.3)_ = 4.5866 minimum for significant comparisons, p <0.011), p<0.023, n_PYR_ = 163 neurons; n_VIP_ = 225 neurons; n_NDNF_ = 128 neurons. (**F-J)** SAM noise (4-128Hz modulation rate, 25-100% modulation depth, 65dB) presented as in (**A-E**). **(G)** Mixed-effects two-way ANOVA, neuron group x octave bin with post-hoc t-test with Bonferroni correction within octave bin compared to PYR (F_24,6621_ = 2.2046, p=0.0006), ^+^VIP or ^Δ^NDNF significantly different from PYR (F_20,4353_ = 3.2203, p<0.0001), ^+^VIP or ^Δ^NDNF significantly different from PYR, (F_2,303.1_ ≤ 3.9509, p <0.02), p<0.04, n_PYR_ = 160 neurons; n_VIP_ = 131 neurons; n_NDNF_ = 123 neurons. **(H)** Mixed-effects one-way ANOVA with Tukey’s post-hoc test, (F_2,44.41_ = 11.3608, p <0.001), **p<0.0001, *p=0.0015, n_PYR_ = 160 neurons; n_VIP_ = 131 neurons; n_NDNF_ = 123 neurons. **i** Mixed-effects two-way ANOVA, neuron group x octave bin with post-hoc t-test with Bonferroni correction within octave bin compared to PYR, (F_20,4493_ = 9.7686, p<0.001), ^+^VIP or ^Δ^NDNF significantly different from PYR, (F_2,270.7_ ≤ 14.541, p <0.001), p<0.001, n_PYR_ = 160 neurons; n_VIP_ =131 neurons; n_NDNF_ = 123 neurons. **(J)** Mixed-effects two-way ANOVA, neuron group x octave bin with post-hoc t-test with Bonferroni correction within octave bin compared to PYR (F_20,4470_ = 12.4477, p<0.001), ^+^VIP or ^Δ^NDNF significantly different from PYR, (F_2,141.7_ ≤ 18.19, p <0.001), p<0.0001, n_PYR_ = 160 neurons; n_VIP_ =131 neurons; n_NDNF_ = 123 neurons.

We also asked whether VIP and NDNF interneurons showed selectivity for specific FM sweeps, which varied in speed and directionality. We observed robust responses to FM stimuli within subsets of all three neuronal subtypes (24.2% PYR, 12.3% VIP, 11.2% NDNF, Fig. 4a). All subtypes demonstrated sweep preferences across the range of speeds and direction (Fig. 4b). We calculated a sweep direction selectivity index to characterize each neuron’s preference for upward versus downward sweeps, and a speed selectivity index to characterize each neuron’s preference for slow versus fast sweep rates. Interestingly, more VIP and NDNF interneurons preferred downward FM sweeps, and more NDNF interneurons preferred slow FM sweeps (Fig. 4c).

**Figure 4.**
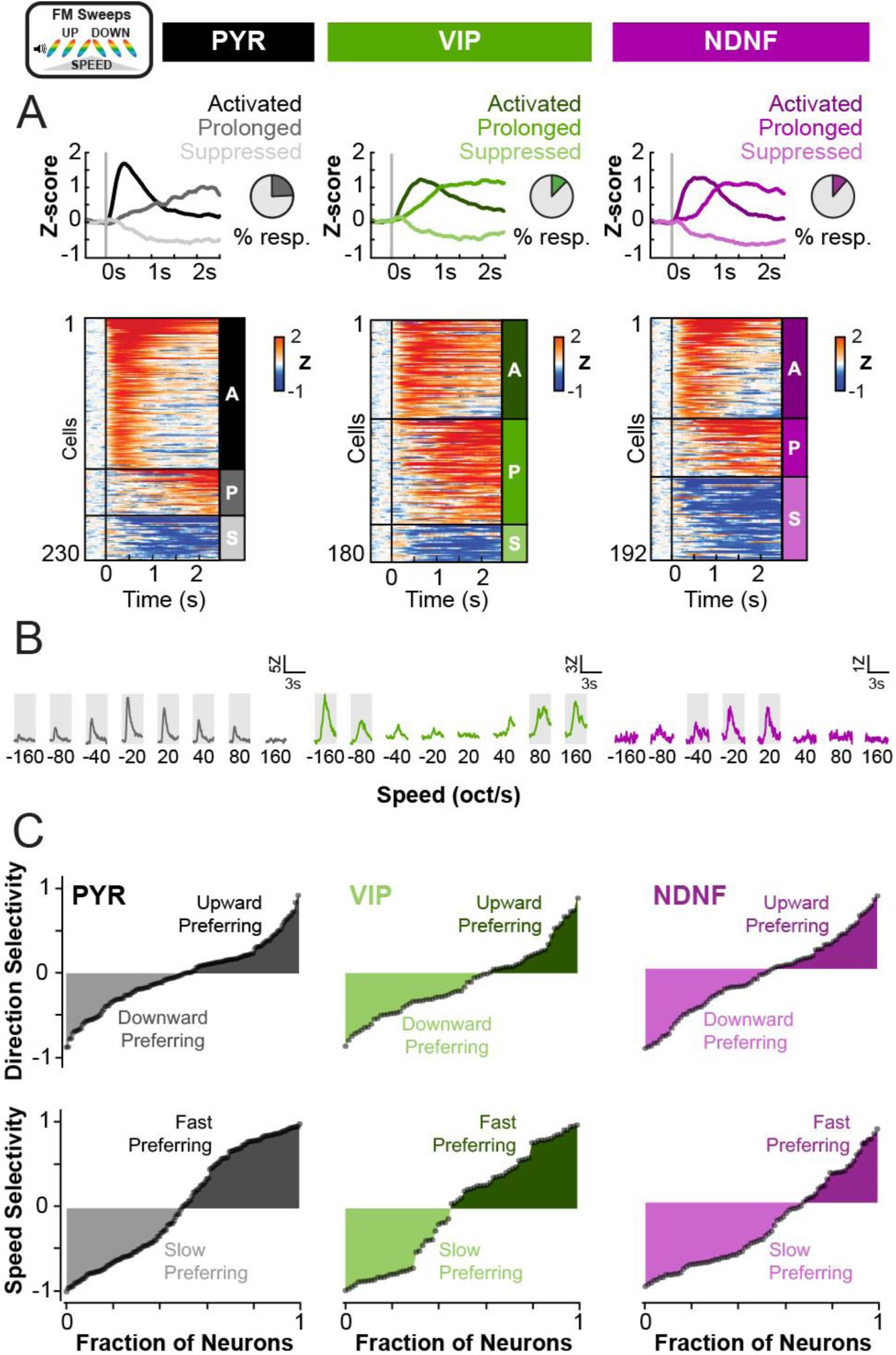
NDNF neurons prefer slow frequency modulated sweep rates. **(A)** For PYR, VIP, and NDNF neuronal populations: top, averaged responses for *activated*, *prolonged*, and *suppressed* neurons to frequency modulated (FM) sweeps (3-96 kHz, 65dB, ±20-160 oct/sec). Vertical line denotes sound onset. Pie chart represents the percentage of total neurons responsive to FM sweep stimuli. Bottom, raster plot showing all mean Z-scored reliable responses to FM stimuli. Each row represents a single neuron. Data are sorted by maximal response within *activated* (A), *prolonged* (P), and *suppressed* (S) profiles. **(B)** Example FM sweep receptive field for a single neuron with reliable responses denoted by gray boxes. **(C)** Sweep direction selectivity index (top row) for *activated* neurons, ordered from downward to upward preferring, and speed selectivity index (bottom row) for *activated* neurons, ordered from slow to fast preferring. Each dot represents a single neuron. n_PYR_ = 144 neurons; n_VIP_ =73 neurons; n_NDNF_ = 80.

Together, these results reveal that subsets of VIP and NDNF interneurons in L1 demonstrate robust responses and clear tuning to a range of complex sounds. However, these populations display large variations in responses to these sounds across trials, particularly in response to non-preferred stimuli. Examining the trial-to-trial reliability of responses, however, unveils ‘reliable’ tuning curves for these interneurons that are narrower than the L2/3 PYR neurons. This finding suggests that VIP and NDNF interneurons may display remarkable selectivity for a narrow range of simple and complex sound stimuli. Further, neurons are most reliable at their preferred stimuli, suggesting that the trial-to-trial variability of sensory-evoked responses may carry information about specific sound stimuli. L1 interneuron populations in particular, may rely on these differences in trial-to-trial variability to encode stimuli rather than response amplitudes alone.

### Locomotor activity modulates the sound-evoked and spontaneous activity of VIP and NDNF interneurons

Our results reveal that low trial-to-trial reliability distinguishes L1 interneurons from PYR neurons. VIP and NDNF interneurons may inherit sound frequency tuning from direct inputs from the primary MGBv, but their tuning may be obscured by low trial-to-trial reliability. What underlies this variability in L1 interneuron responses? L1 interneuron activity may be modulated by inputs from diverse brain regions that convey information about behavioral state, including locomotion^9,13,36^. We therefore examined how running impacts response amplitude and reliability of VIP and NDNF interneurons. Consistent with previous reports on the effects of locomotion on PYR neuron activity^53,54,55^, we observed a reduction in the peak amplitude of *activated* or *prolonged* PYR neurons and a reduction in the trough amplitude of *suppressed* PYR neurons during motor bouts (Fig. 5a-d). This reduction in response amplitude was associated with a decrease in trial-to-trial reliability (R_Sound_) while running. In contrast, VIP and NDNF interneurons exhibited an increase in response amplitude, and NDNF interneurons showed an increase in response reliability during running (Fig. 5a-d).

**Figure 5.**
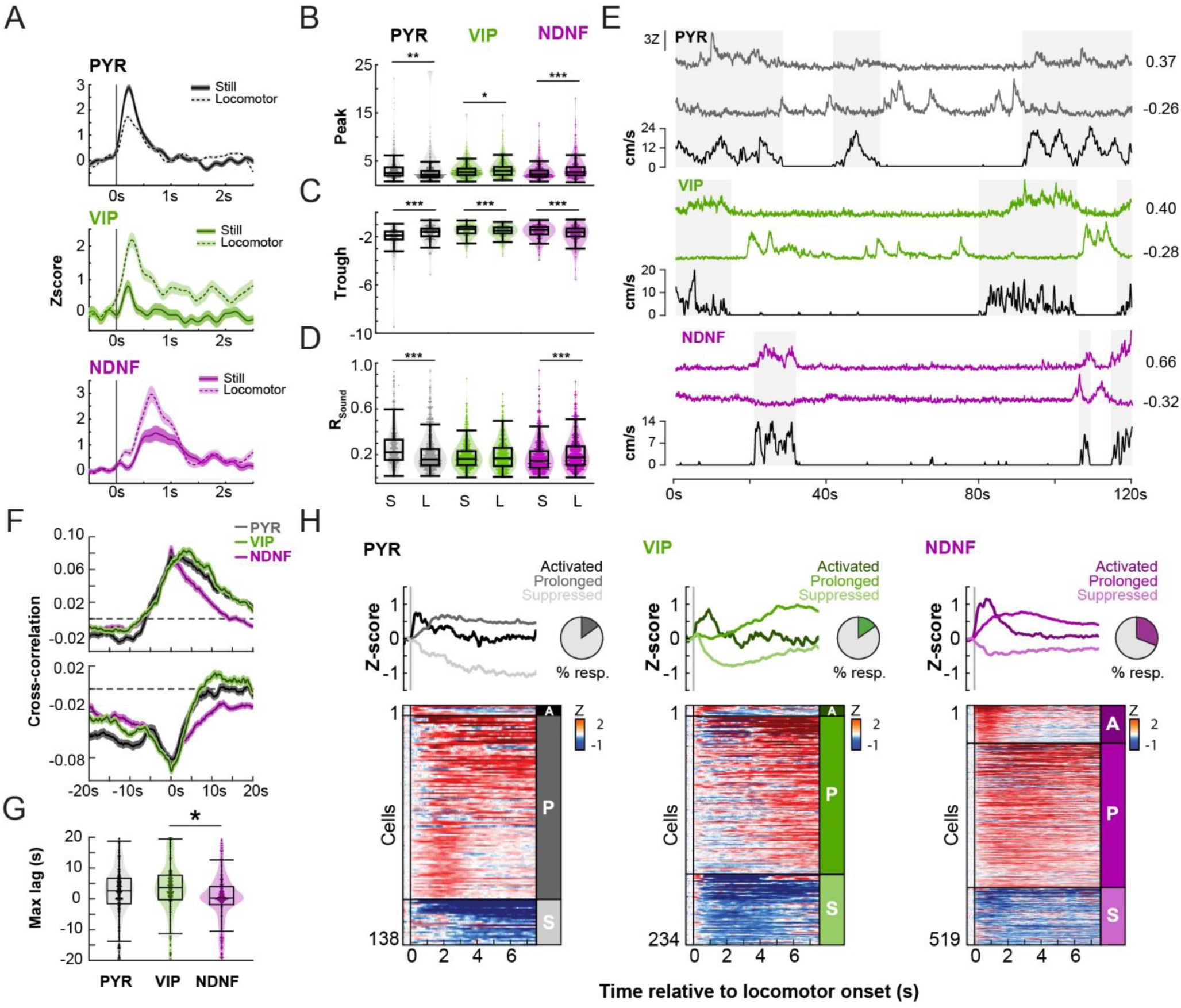
Locomotor activity enhances the amplitude and reliability of sound-evoked responses in NDNF neurons. **(A)** Averaged responses to noiseburst stimuli recorded from example PYR, VIP, and NDNF neurons during stillness (S) and locomotor activity (L). Shading shows +/- standard error. **(B)** Peak response from *activated* and *prolonged* neurons during stillness and locomotion. Mixed-effects two-way ANOVA, neuron group x running status with Tukey’s post-hoc test, (F_2,3645_ =22.8552, p<0.001), ***p<0.001, **p = 0.004, *p=0.028, n_PYR_ = 557 neurons; n_VIP_ =690 neurons; n_NDNF_ = 656 neurons. **(C)** Trough responses from *suppressed* neurons. Mixed-effects two-way ANOVA, neuron group x running status with Tukey’s post-hoc test, (F_2,1583_ =24.1828, p<0.001), ***p<0.001, n_PYR_ = 275 neurons; n_VIP_ =336 neurons; n_NDNF_ = 459 neurons. **(D)** Response reliability during stillness and running. Mixed-effects two-way ANOVA, neuron group x running status with Tukey’s post-hoc test, (F_2,5165_ = 59.0422, p<0.001), ***p<0.001, **p = 0.004, *p=0.028, n_PYR_ = 742 neurons; n_VIP_ = 921 neurons; n_NDNF_ = 1003 neurons. **(E)** Example traces from PYR, VIP and NDNF neurons that are positively (top) or negatively (middle) correlated with locomotor activity (bottom trace). Number denotes the cross-correlation coefficient of each neuron with locomotor activity at zero lag. Recordings were made during periods of spontaneous activity with no sound presentation. **(F)** Average cross-correlograms between spontaneous neural activity and locomotor activity for each population. Top, Neurons positively correlated with locomotor activity. Bottom, Neurons negatively correlated with locomotor activity. **(G)** Lag time of peak cross correlation for neurons positively correlated with locomotor activity. Mixed-effects one-way ANOVA with Tukey’s post-hoc test, (F_2,26.54_ = 5.0382, p=0.014), *p=0.01, n_PYR_ = 951 neurons; n_VIP_ = 1197 neurons; n_NDNF_ = 988 neurons. **(H)** For PYR, VIP, and NDNF neuronal populations: top, averaged responses for *activated*, *prolonged* and *suppressed* neurons to the onset of locomotor bouts. Vertical line denotes bout onset. Pie chart represents the percentage of total neurons responsive to locomotion onset. Bottom, raster depicting all mean Z-scored neuron responses to locomotor bouts. Each row represents a single neuron. Data are sorted by maximal response within *activated* (A), *prolonged* (P), and *suppressed* (S) profiles.

We also asked how locomotion impacts the activity of L1 interneuron populations in the absence of sound. NDNF interneurons showed the strongest positive, time-locked correlations with locomotor activity. Spontaneous NDNF activity was tightly coupled with locomotion while VIP and PYR activity lagged the locomotor trace (Fig. 5e-g). Furthermore, 31% of NDNF interneurons responded to the onset of a motor bout compared to 13% of VIP and 14.8% of PYR neurons (Fig. 5h). This strong modulation of NDNF interneurons by locomotion may underlie the increased amplitude and reliability of sound-evoked responses within this interneuron population observed during running. Together, these results indicate that fluctuations in L1 interneuron activity, particularly NDNF interneurons, may arise from inputs conveying information about specific behavioral states.

### L1 interneuron ensembles share trial-to-trial variability

Trial-to-trial variability in sensory-evoked responses can be cell autonomous, resulting from variable synaptic and intrinsic cellular properties that lead to fluctuations in synaptic transmission or dynamic spike thresholds^56^. Response variability can also be shared across neuronal networks^57^, reflecting shared non-sensory inputs^58^ or direct coupling between neighboring neurons^59^. In addition to receiving diverse non-auditory inputs^14^, interneurons within L1 form complex networks that are interconnected by inhibitory synapses and gap junctions^33,60,61^. We therefore asked whether the trial-to-trial variability we observed is shared among L1 interneurons. We measured pairwise signal and noise correlations among neurons within the same field-of-view. Signal correlation quantifies the similarity of neuronal responses to repeated trials of the same sound stimulus whereas noise correlation quantifies the stimulus-independent trial-to-trial variability that may arise from common, non-auditory inputs or interactions between neurons. As expected from previous work^48^, signal and noise correlations were stronger between PYR neurons that were in close spatial proximity (Supplementary Fig. 2). We also observed this relationship in VIP and NDNF interneurons (Supplementary Fig. 2) despite the greater inter-neuronal distance that characterizes these sparse populations. To account for differences in inter-neuronal distance between the three populations, we used distance as a covariate in subsequent correlation analyses. Interestingly, all three populations exhibited a similar percentage of pairs showing significant signal correlations (36% PYR, 46% VIP, 31.6% NDNF, Fig. 6a). However, we observed higher signal correlations for PYR neurons as compared to VIP and NDNF interneurons, consistent with previous reports that ensembles of PYR neurons show co-tuning to sound stimuli^47,49^ (Fig 6a,b). However, we found that both VIP and NDNF interneurons showed significantly higher positive noise correlations as compared to PYR neurons. Finally, NDNF interneurons showed significantly stronger negative noise correlations than PYR and VIP neurons (Fig 6c,d), consistent with strong inhibitory synaptic coupling between NDNF interneurons^61^. Together, these results demonstrate that the VIP and NDNF interneuron populations display greater shared trial-to-trial variability than PYR neurons, suggesting that strong interactions between neighboring L1 interneurons or common non-auditory inputs may contribute to their reduced sound-evoked response reliability.

**Figure 6.**
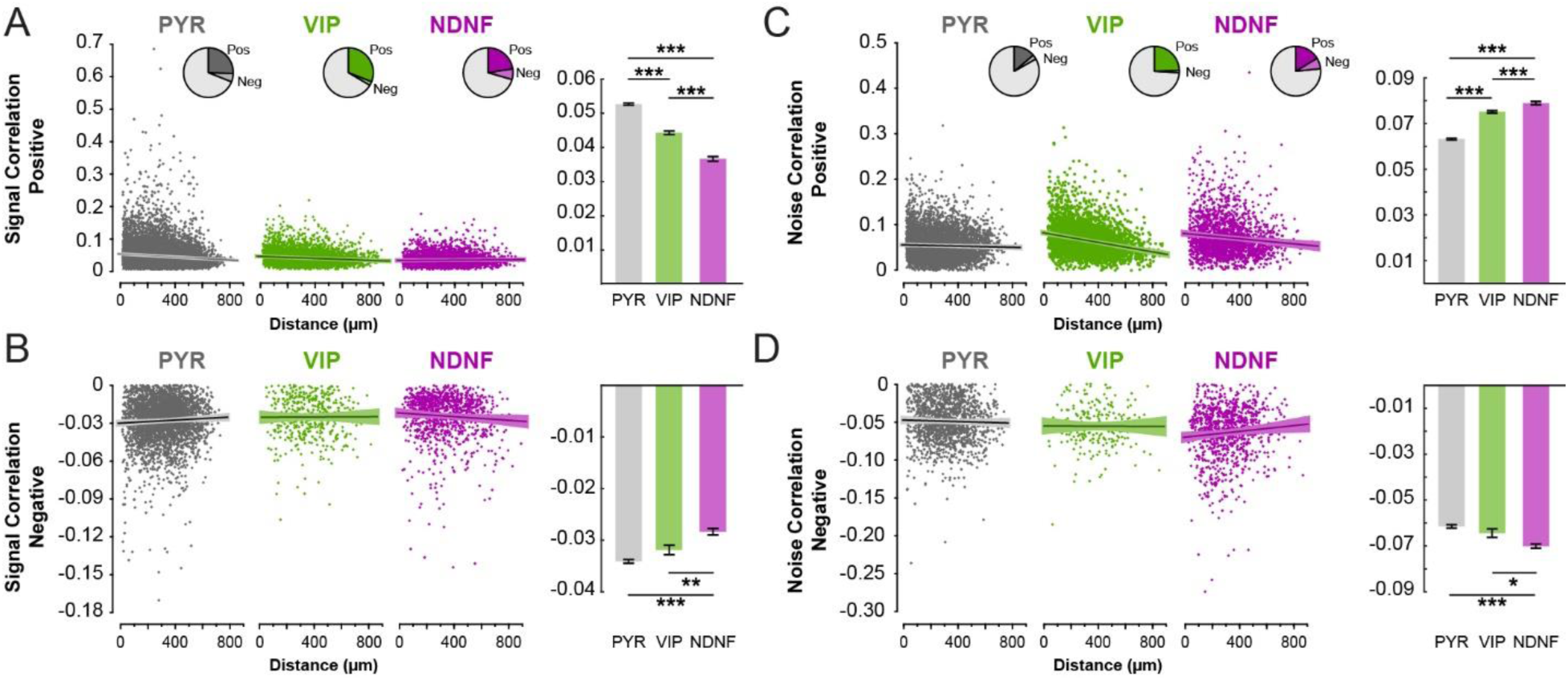
VIP and NDNF interneurons show weaker signal correlations but stronger noise correlations than PYR neurons. **(A)** Positive signal correlations between pairs of PYR, VIP, and NDNF neurons. Left, positive signal correlation as a function of spatial distance between neuron pairs overlaid by linear fit. Single points denote mean correlation value across auditory stimuli for each neuronal pair. Pie chart represents the percentage of total neurons with significantly positive and negative signal correlations. Right, mean of positive signal correlations. Mixed-effects one-way ANOVA with Tukey’s post hoc test, (F_2,26752_ = 258.5059, p<0.001), ***p<0.0001, n_PYR_ = 14861 neuron pairs; n_VIP_ = 5141 neuron pairs; n_NDNF_ = 3268 neuron pairs. **(B)** Negative signal correlations depicted as in (**A**). Mixed-effects one-way ANOVA with Tukey’s post hoc test, (F_2,5015_ = 32.3813, p<0.001), ***p<0.0001, **p=0.004, n_PYR_ = 3329 neuron pairs; n_VIP_ = 492 neuron pairs; n_NDNF_ = 1061 neuron pairs. **(C)** Pairwise positive noise correlations depicted as in (a). Mixed-effects one-way ANOVA with Tukey’s post hoc test, (F_2,12829_ = 262.158, p<0.001), ***p<0.0001, n_PYR_ = 6653 neuron pairs; n_VIP_ = 3625 neuron pairs; n_NDNF_ = 2169 neuron pairs. **(D)** Negative noise correlations depicted as in (**A**). Mixed-effects one-way ANOVA with Tukey’s post hoc test, (F_2,2639_ =26.1381, p<0.001), ***p<0.001, *p=0.019, n_PYR_ = 1436 neuron pairs; n_VIP_ = 241 neuron pairs; n_NDNF_ = 944 neuron pairs.

### VIP and NDNF interneuron activity is significantly modulated by non-auditory sources

Robust modulation of L1 interneurons by behavioral states, such as locomotor activity (Fig. 5), and shared activity between neighboring L1 interneurons (Fig. 6) suggest that features beyond bottom-up auditory inputs influence their activity. We therefore predicted that factors other than auditory stimuli are necessary to model the activity of these interneurons. We implemented a generalized linear model (GLM) to model the activity of individual VIP, NDNF, and PYR neurons. We quantified the prediction performance of each model as the correlation between the predicted and actual neuron’s activity on a subset of trials not used for fitting. Relying on sound-related stimuli alone to model single-cell L1 interneuron activity led to poor predictions as compared to PYR neurons (Fig.7a-c). However, when we incorporated non-sensory factors as variables, such as locomotor activity or the responses from neighboring neurons, we were able to capture substantial variance across trials and our models performed similarly across all three cell classes (Fig. 7 d-f). These results suggest that changes in the mouse’s behavioral state or interactions between neighboring neurons may contribute significantly to the trial-to-trial variability observed in L1 interneuron populations.

**Figure 7.**
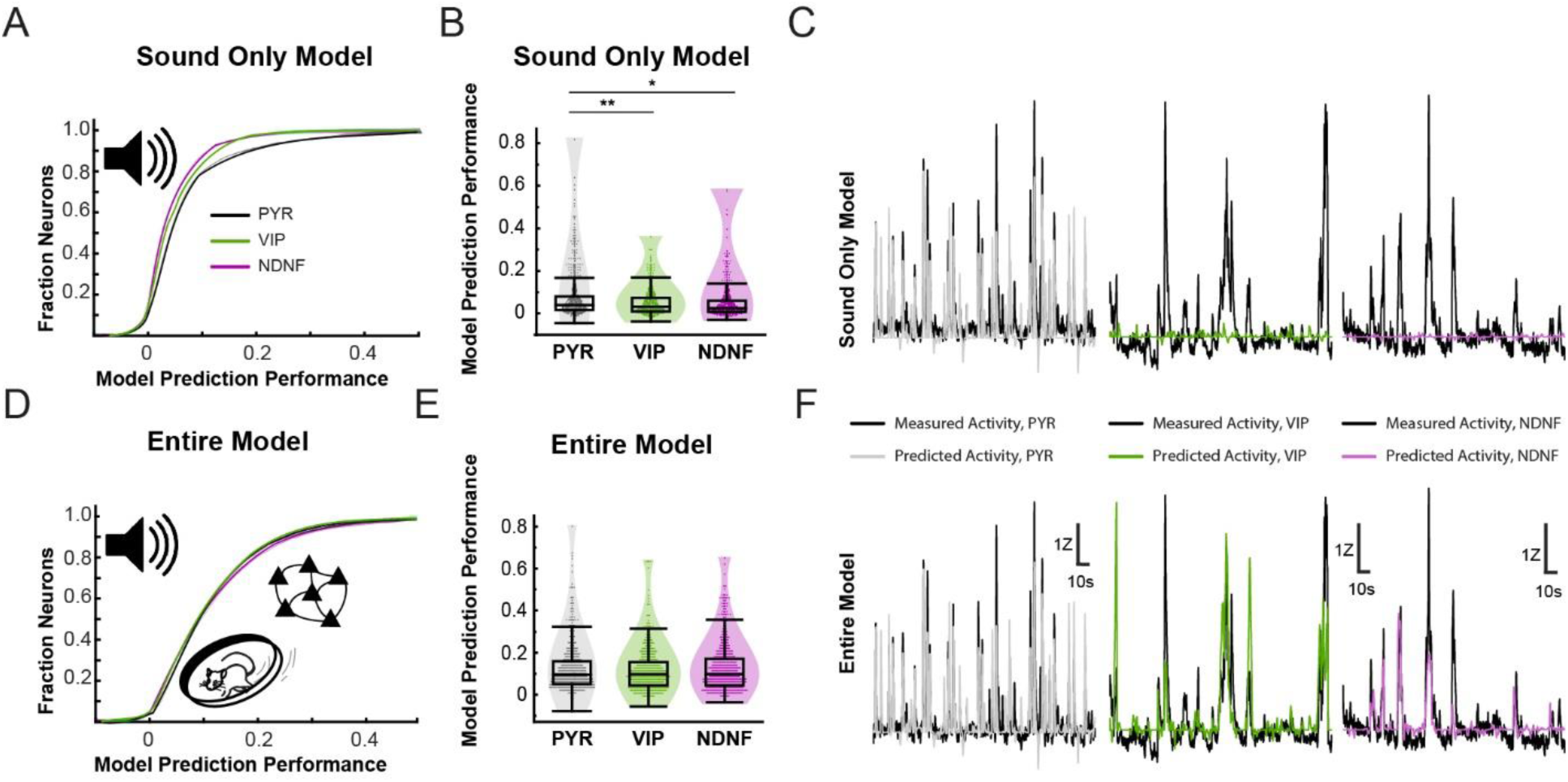
VIP and NDNF interneurons are significantly modulated by non-auditory sources. Generalized linear model (GLM) of PYR, VIP and NDNF responses. A GLM was fit to training data to predict the calcium responses of individual neurons using either sound stimulus predictors alone (**A-C**; sound only model) or sound stimulus, neighboring neuron activity, and locomotor activity predictors (**D-F**; entire model). **(A)** Cumulative distribution function of sound only model prediction performance for all neurons, calculated as the correlation coefficient of measured versus predicted traces. **(B)** Mean prediction performance of sound only model. Mixed-effects one-way ANOVA with Tukey’s post-hoc test, (F_2,21.91_ = 6.4974, p = 0.005), **p=0.009, *p =0.01, n_PYR_ = 687 neurons, n_VIP_ = 758 neurons, n_NDNF_ = 816 neurons. **(C)** Example PYR, VIP and NDNF neuron traces (black) and predicted fits with sound only GLM model (gray/green/purple). **(D)** Cumulative distribution function of entire model prediction performance as in (**A**). **(E)** Mean prediction performance of entire model. Mixed-effects one-way ANOVA (F_2,21.49_ = 1.1311, p =0.028), n_PYR_ = 687 neurons, n_VIP_ = 758 neurons, n_NDNF_ = 816 neurons. **(F)** Example PYR, VIP and NDNF neuron traces (black) and predicted fits with entire GLM model (gray/green/purple).

## Discussion

L1 of sensory cortex is a site of convergence for long-range projections originating from diverse brain regions that relay both sensory information and internal signals such as behavioral states and expectations^16,17^. The GABAergic inhibitory interneurons that populate L1 receive these diverse inputs^17^ and are therefore uniquely positioned to process sensory information in an experience- and state-dependent manner. Understanding whether and how the activity of L1 interneurons represent discrete features of the sensory environment in awake, behaving animals will provide important insights into their contributions to sensory perception and learning. Here, we found that both simple and complex sounds evoke robust calcium signals in VIP and NDNF interneurons within L1 of ACtx. Surprisingly, both interneuron subtypes display selective responses to specific spectrotemporal features of sounds, similar to the well-studied L2/3 PYR neurons (Figs. 2-4). However, one notable difference between these neuronal populations is that the sound-evoked responses of L1 interneurons are less reliable, exhibiting high trial-to-trial variability (Fig. 1). We discovered that this variability is modulated by behavioral state (Fig. 5) and shared across neighboring L1 interneurons (Fig. 6). L1 interneurons have recently emerged as ‘master regulators’ that receive internal signals about behavioral states and past experiences to gate cortical activity and plasticity^16,17^. Our study importantly adds that these interneurons are also key sites of sensory processing that may exert their effects on cortex in both a stimulus- and state-dependent manner. However, the mechanisms by which these interneurons process sensory information, utilizing trial-to-trial response variability to carry information about the stimulus and behavioral cues, may be fundamentally different than the well-studied PYR neurons.

Information about specific features of the sensory environment is transferred to cortex via axons from the primary thalamic nuclei. We recently reported that the majority of monosynaptic thalamic inputs to VIP and NDNF interneurons within L1 of the ACtx arises from the primary auditory thalamus, the MGBv^33^. This connectivity predicts that VIP and NDNF interneurons are not only sound responsive, but also inherit tuning properties from the frequency-selective MGBv neurons. Indeed, we found that L1 NDNF and VIP interneurons exhibit robust calcium responses to sounds. This finding is consistent with other recent studies showing that sound stimuli elicit responses in NDNF^14^ and VIP^6,37,38^ interneurons during auditory-driven behaviors. Moreover, we showed that subsets of VIP and NDNF interneurons show narrow tuning to the frequency of pure tone stimuli (Fig. 2), as well as tuning to spectrotemporal features of complex amplitude- and frequency-modulated sounds (Figs. 3 and 4). FM sweeps are key features of vocalizations^62,63,64^, and amplitude-modulated sounds mirror envelope features of naturalistic noises^63,65^. Together, these studies suggest that L1 interneurons may contribute to the decoding of physiologically relevant sounds.

While L1 interneurons and PYR neurons exhibited surprising similarities in sound-evoked responses and tuning, one striking difference among these neuronal populations is their trial-to-trial variability. Whereas subsets of PYR neurons showed similar responses across trials of matched sound stimuli, most L1 interneurons exhibited large fluctuations in response amplitude across trials (Fig. 1). This is consistent with the all-or-none sensory evoked-responses that have been found for L1 interneurons using voltage imaging in the whisker cortex^61^. These all-or-none responses may be unique to L1 interneurons, resulting from winner-takes-all network dynamics within L1^61^ and/or from specialized biophysical properties that produce persistent firing modes^66^. This variability led to noisy receptive fields that are consistent with previous *in vivo* measurements of cortical interneurons^45,67^. Throughout our analyses, we leveraged reliability measures as a tool to highlight robust tuning properties within a seemingly noisy receptive field.

Trial-to-trial variability is expected in neuronal populations given stochastic synaptic transmission and dynamic spike thresholds^61^. However, recent studies have highlighted that trial-to-trial variability can reflect more than just random noise, but instead represent network dynamics that contribute to the decoding of stimulus-related information^68,69,70^. For example, the trial-to-trial variability of neurons within the rat whisker thalamus contributes information about stimulus location^71^. We found that the trial-to-trial variability observed in L1 interneurons also carried information about the sound stimulus (Figs. 2 and 3). L1 interneurons displayed low reliability to non-preferred sound stimuli, leading to just a narrow range of stimuli that elicited reliable responses. When focusing on these reliable responses, L1 interneurons therefore exhibited narrow tuning curves. These findings point to the interesting possibility that subsets of inhibitory interneurons, while traditionally thought of as poorly tuned^67^, may in fact be remarkably selective to specific features of the sensory environment.

Our findings suggest that L1 interneurons convey information about the spectrotemporal features of sounds, but that the coding strategy of these interneurons may differ from the well-studied PYR neurons by relying on trial-to-trial variability. Along the ascending sensory systems, there is a transition from neurons that exhibit little noise in the early stages of sensory processing to neurons that show elevated noise levels at higher processing areas^72^. This change in the degree of variability across the sensory hierarchy is thought to represent a progression in the population coding strategy, from rate coding to ‘sparse’, efficient coding. ‘Sparse’ coding minimizes the occurrence of metabolically expensive action potentials, increasing computational and metabolic efficiency^73^. High trial-to-trial variability therefore is thought to be an emergent feature of higher processing regions that integrate diverse inputs to reduce the redundancy of information across neurons. Moreover, theoretical work predicts that this variability is exploited in the brain to generate error signals that are necessary for memory storage during associative learning^74^. This may be particularly important for L1 interneurons that process inputs from diverse brain regions^13,14,33^ and undergo robust plasticity during development^15^ and adult associative learning^14^. Our findings thus motivate future experimental and theoretical work to understand how the trial-to-trial variability of L1 interneurons supports moment-to-moment auditory perception and learning.

Recent studies have suggested that periods of trial-to-trial variability can reflect specific brain states that support efficient information processing^68,69,70,75,76^. What modulates the variability of L1 interneurons? Here we found that a change in behavioral state reduced variability of sound-evoked responses in NDNF interneurons (Fig. 5). Variations in behavioral state, including locomotor bouts, are tightly correlated with states of high arousal, as measured by fluctuations in pupil diameter^54^. Previous work in visual cortex reports that NDNF interneuron activity correlates with arousal states^77^, which serves as a mechanism to gain-modulate PYR neurons^13^. While running bouts enhanced the response reliability of NDNF interneurons and increased the response amplitude of NDNF and VIP interneurons, we did not observe a change in VIP response reliability during running. Future studies that identify specific behavioral states that modulate response reliability in VIP interneurons will prove interesting in uncovering the role of distinct long-range influences on these two different L1 subpopulations. Together, these findings motivate future work investigating how unique behavioral states modulate the reliable encoding of sensory signals within L1 interneurons.

In addition to long-range, motor input onto L1 interneurons, local network dynamics may drive variability within the VIP and NDNF interneuron populations. These dynamics may reflect complex interactions between L1 interneurons that are known to be interconnected by gap junctions^60^, inhibitory synapses^33,61,66^, and excitatory cholinergic synapses^78,79,80^. Consistent with this prediction, we found that VIP and NDNF interneurons showed significantly greater positive and negative noise correlations as compared to PYR neurons (Fig. 6). Greater positive correlations may be driven by common long-range excitatory connections, such as motor inputs, or may result from direct electrical^60^ or excitatory synaptic connections^78,79,80^ between L1 interneurons, while stronger negative noise correlations are consistent with the robust inhibitory synaptic connections among L1 interneurons^33,61,66^. Indeed, we find that L1 interneuron activity relies on the aggregation of multiple sources of input (Fig. 7).

Our findings highlight the function of L1 interneuron populations in the encoding of specific features of complex sensory environments. While previous studies have focused on responses of VIP and NDNF interneurons to non-sensory and contextual cues such as reinforcing stimuli and learned associations^14,37,38,39^, our results emphasize that these interneurons are indeed sensory neurons that exhibit narrow tuning to distinct stimuli. Mapping the functional circuitry of L1 interneurons will be critical for understanding how this sensory information is transmitted to specific cortical targets to underlie state-dependent modulation and plasticity. The interneurons within superficial cortex send spatially-precise outputs^14,15,66,81^, some of which descend vertically to deeper-layer inhibitory interneurons to create ‘holes’ in inhibitory networks^82^ and gain-modulate cortical columns^13,83^. Our findings support a new model that the modulation of subnetworks of co-tuned L1 interneurons, rather than blanket L1 interneuron activation by global contextual cues, is critical for moment-to-moment sensory perception and learning. Future studies may thus highlight that modulating the reliability of small ensembles of co-tuned L1 interneurons is a mechanism for filtering sensory information and relaying contextual cues within complex cortical circuits.

## Data availability

Raw and summary data that support the findings of this study are available from the corresponding author upon request.

## Code availability

Source code is available on Github (https://github.com/takesian-lab/Sweeney-Thomas-et-al).

## Acknowledgments

This work was supported by funds from the NIH NIDCD R01DC018353, and NIDCD F32DC018211 (to C.G.S.), the Nancy Lurie Marks Family Foundation, the Bertarelli Foundation, the Pew Latin American Fellows Program (to L.G.V.), the Natural Sciences and Engineering Research Council of Canada (to M.E.T.), and the Fonds de recherche du Québec – Nature et technologies (to M.E.T.). We thank Dr. Daniel Polley (Mass Eye and Ear, Harvard Medical School) for valuable feedback and advice, Dr. Polley’s lab members for technical assistance and discussions, Ken Hancock for software support, Rahul Brito and Ian Griffith for analytical support, and Cathryn MacGregor and Benjamin Glickman for animal care.

## Author contributions

Conceptualization, CGS, MET, and AET; Investigation, CGS, MET, KES,and LGT.

Software and analysis, CGS, MET, and AET.; Visualization, CGS, MET, and AET; Writing, CGS and MET, Review & Editing, CGS, MET, AET, and LGV.

## STAR Methods

### Mice

VIP-cre (JAX, 010908) and NDNF-cre (JAX, 028536) mouse lines were crossed with the Ai148 line (JAX, 030328) to selectively express the calcium indicator, GCaMP6f, in VIP and NDNF interneuron populations. To selectively express GCaMP6f in pyramidal neurons (PYR), Ai148-negative mice from both strains were injected with AAV9-CaMKII-GCaMP6f-WPRE-SV40 (2.7×10^12 gc/ml, Addgene, 100834). The number of mice per group was PYR N = 6 (2 females), VIP N = 11 (6 females), and NDNF N = 10 (4 females). Experiments were carried out in adult mice aged >P60. Mice were housed in groups of at least two with food ad libitum and a running wheel provided for environmental enrichment. A 12:12 hour light:dark cycle was maintained with experiments performed during the light period. All experimental and surgical procedures were approved by the Mass Eye and Ear Animal Care Committee.

### Cranial window surgeries and viral injections

Mice were implanted with a cranial window and headplate to allow for chronic, head-fixed imaging of awake mice. Prior to surgery, mice were anesthetized with isoflurane (5% induction, 1–2% maintenance) and administered buprenex (0.5mg/kg, s.c.) and meloxicam (2mg/kg, s.c.). Petroleum ophthalmic ointment was applied to the eyes and body temperature was maintained with a heating pad. After removing hair from the surgical area, the skin was cleaned by alternating scrubbing with 70% ethanol and betadine three times before an incision was made over the midline. A titanium or stainless steel headplate (1.5 cm x 1.7 cm x 0.1 cm, L x W x thickness; < 1g) was affixed to the skull using dental cement (Patterson Dental, S380). A 3mm circular craniotomy was drilled over the left temporal ridge directly anterior to the lambdoid suture. For virally-injected mice, 200nl of virus (AAV9-CaMKII-GCaMP6f-WPRE-SV40, 1:10) was injected at a depth of 200µm below the pial surface and a rate of 25nl/min via stereotaxic microinjection using a glass pipette controlled by a syringe (Hamilton) pump (WPI), which was left in place for an additional 5 minutes following each injection. Injections were performed in 2-3 locations approximately 2.3-2.7mm posterior to bregma and parallel to the temporal ridge avoiding vasculature. Finally, a cranial window consisting of two 3mm and one 4mm round microscopy glass coverslips glued together with a UV-cured optical adhesive (Norland, 7106) was implanted. Silicone adhesive (WPI, KWIK-SIL) was used to create a seal between the skull and window. The coverslip was affixed to the surrounding skull with dental cement. Meloxicam was administered for post-operative analgesia 24 and 48 hours after surgery. Following cranial window implantation, mice recovered for at least five days before imaging.

### Two-photon calcium imaging

Cellular two-photon calcium imaging was performed in awake mice using a Bruker Ultima multiphoton imaging system in a sound-attenuating light-tight enclosure supported on an optical floating table (Newport). The entire cranial window was imaged using a widefield epifluorescence microscope at both 4x (Olympus, 0.13NA) and 16x (Nikon, 0.8NA, water-immersion) magnification. Widefield illumination was achieved with a 120W mercury arc lamp (Lumen, XCite 120Q) and filter cube (Filters ET470/40x and ET525/50m; dichroic T495LPXR, Chroma) and captured at 20Hz with a CMOS camera (Hamamatsu, ORCAp-Flash4.0 LT+). Two-photon imaging was performed with the same 16x objective and an 8kHz resonant galvanometer scanner for high-speed imaging (30Hz, 512 x 512 pixels, 827 x 827 µm). A motorized nosepiece allowed the imaging system to rotate around the head of the mouse, permitting the lens to be positioned at an angle of ∼55° to image the auditory cortex. Laser light was delivered from a tunable, ultrafast laser (InSight X3, Spectraphysics) tuned to 920nm at approximately 20-40mW power measured at the objective and adjusted with Pockels cells (Conoptics, 302RM). Emission signals were separated by a 700nm shortpass dichroic mirror (Hunt Optics and Imaging) and bandpass filtered before detection was performed with high-sensitivity large-angle GaAsP photomultiplier modules (PMTs, Hamamatsu, model 10770). For each mouse, one to four non-overlapping 2P fields-of-view (FOVs) were imaged to obtain responses from multiple regions within the auditory cortex. The imaging depth, defined as the distance from the pial surface, was chosen per mouse to obtain FOVs in superficial cortex with a high density of cells (mean depth: L2/3 PYR 219µm, N=6 FOVs; L1/2 VIP 168µm, N=19 FOVs; L1 NDNF 87µm; N=20 FOVs). During all imaging sessions, mice were head-restrained and allowed to run on a wheel designed to prevent movement-related sound. The speed of voluntary running was monitored using a custom tachometer and recorded with a sampling rate of 10Hz during image acquisition.

### Acoustic stimuli

Mice were passively presented with acoustic stimuli generated and delivered using custom hardware and software designed for auditory behaviors by the Eaton-Peabody Laboratory engineers (LabView). The acoustic stimuli were delivered through a calibrated free-field speaker positioned 7cm from the mouse’s right ear. The stimuli (all including 5ms cosine on/off ramps) were broadband white noise bursts (75dB, 200ms), pure tones (4-32kHz, 20-80dB, 200ms), frequency-modulated (FM) sweeps (3-96kHz, +/-20-160 octaves/s, 31-250ms, 65dB), sinusoidal amplitude-modulated white noise (broadband, 4-128Hz modulation rate, 0-100% modulation depth, 500ms, 65dB), and sinusoidal amplitude-modulated pure tones (4-32kHz, 10Hz modulation rate, 0-100% modulation depth, 500ms, 65dB). Each stimulus was repeated for 5-20 trials and played in a pseudo-random order with inter-stimulus intervals of 5.5s. Blank trials, in which no sound was presented, were randomly interleaved with sound stimulus trials at a rate of 10-15%. Acoustic stimuli were recorded from one FOV per day per mouse.

### Anatomical verification of imaging positions

To accurately determine the position of FOVs within the ACtx, dyes were injected in each recording site following the completion of imaging. Mice were anesthetized with isoflurane (5% induction, 1–2% maintenance) and placed in a stereotaxic surgical frame. The cranial window was removed by drilling the dental cement holding it in place and gently lifting the window with forceps. The vasculature patterns from widefield images were then matched to the cortical surface. A glass micropipette dipped in either DiI or DiD paste (ThermoFisher, N22884) was inserted at a depth of 400µm below the cortical surface at each imaging site. Mice were injected with fatal plus (1.8 mg/kg, i.p.), and intracardiac perfusion with paraformaldehyde was performed immediately after dye injection. Once a deep plane of anesthesia was achieved, a midline incision was made using scissors to open the thoracic cavity. A 30-gauge needle attached to silicon tubing was inserted into the left ventricle of the heart to begin perfusion of 10-20ml of saline solution followed by 10-20ml of 0.1M phosphate buffered saline (PBS) containing 4% paraformaldehyde (PFA) at a rate of 5 ml/minute. Flow rate was controlled with a peristaltic pump. After perfusion, the animal was decapitated and the brain was removed and placed in 4% PFA overnight before being transferred to PBS. Thalamocortical slices (80µm)^84^ were cut on a vibratome and stored in PBS-azide. Slices containing dyes were mounted and imaged and auditory cortical regions were determined to be primary, ventral, or dorsal based on anatomical landmarks within the thalamocortical slice.

### Analysis of calcium imaging data

#### Preprocessing

Images were processed using the open-source calcium imaging data analysis pipeline Suite2P^85^ for image registration, x/y motion correction, detection and curation of neurons via cell segmentation and estimation of neuropil activity. Raw fluorescence traces from each cell were corrected for neuropil contamination by subtraction: F_cell corrected_ = F_cell_ - 0.7*F_neuropil_.

#### Detection of stimulus-responsive and reliable neurons

To identify stimulus-responsive neurons, we developed a routine to automatically detect neurons that displayed robust and reliable event-related responses. A trial was defined as the 2.5s period following stimulus presentation and each trial was Z-scored to the 0.5s baseline immediately preceding the sound. Trials of the same stimulus type were averaged together to detect sound-evoked responses. A Z-score threshold of +/-1 was used to detect positive or negative deflections. If a response exceeding this threshold was detected, the response was defined as the portion of the trace containing the maximum or minimum value without changing sign of response. To be considered an event-related response, the minimum response duration was 0.2s and the minimum area-under-the-curve (AUC) was 5 Z-scores. To measure reliability of the detected responses, we computed the average pairwise Pearson’s correlation, R, between each pair of trials and compared these values to the distribution of R values computed from an equal number of blank trials 1000 times via bootstrapping. If the stimulus R value was significantly greater than the blank distribution (one-tailed Z-test, alpha 0.05), the response was deemed to be reliable. Responses with negative R values were not considered. To control for multiple comparisons, the alpha level was adjusted by dividing alpha by the number of responses tested for that neuron. Neurons needed to meet these criteria for at least one stimulus to be included in analyses. Finally, we employed the assumption that neurons reliable to a given stimulus are likely to also respond to sounds within the neighboring stimulus space, such as two tones of similar frequencies. If a neuron was deemed to have at least one reliable response, we used an additional metric to determine whether responses to neighboring stimuli were reliable. We computed the average pairwise Pearson’s correlation, R_neighbor_, between all trials for the reliable stimulus and each of its neighbors. If R_neighbor_ was significantly different from the distribution of R values obtained from circle-shifting the neighboring responses 1000 times (two-tailed Z-test, alpha 0.05), the responses were deemed reliable. Positive and negative R_neighbor_ values were considered, as cells could exhibit positive or negative responses to neighboring stimuli. We controlled for multiple comparisons by dividing the alpha level by the number of neighbors tested.

#### Classification of response types

After classifying the reliability of each cell’s response to every stimulus, average traces were generated from reliable stimuli only. These final traces were subsequently characterized using an automatic routine as either *activated* (having a positive short-latency response), *prolonged* (having a positive response with a peak latency greater than 1s) or *suppressed* (having a negative response). If traces contained both positive- and negative-going components, the trace was defined by the component with the greater AUC.

#### Quantification of receptive field properties and tuning

We performed an automatic process to characterize tuning properties for neurons with activated response types. We defined the best stimulus as the one that evoked the strongest reliable response measured by AUC. To measure tuning strength, d-prime (d′) was computed as in Romero et al.^47^. We computed the average response AUC of the best stimulus and its four neighboring stimuli. We then computed the d′ sensitivity index between this value and the average AUC of five other stimuli selected at random and repeated this process 1000 times. The average of these 1000 repetitions was defined as d′. Normalized receptive fields were generated by shifting the entire receptive field such that the best stimulus was the center column of the receptive field. The remaining area was padded to account for neurons with best stimuli at the limits of stimuli measured.

#### FM Sweep Selectivity Indices

We defined each FM sweep stimuli as upward/downward (</> 0 octaves/s) or slow/fast (</> 50 octaves/s). The FM sweep direction and speed selectivity indices were then computed as Selectivity Index = (Resp_A_ - Resp_B_)/(Resp_A_ + Resp_B_), where Resp_A_ and Resp_B_ are the average AUC for the responses to each category (upward/downward or slow/fast). For neurons with suppressed response types, the sign of each trace was first inverted such that the most suppressed responses were deemed to be the most strongly modulated.

#### Signal and noise correlation analysis

Signal and noise correlations were computed between all sound-responsive cell pairs within each FOV for each stimulus separately. For each pair of cells, we computed correlations during responses to stimuli for which either cell’s responses were deemed reliable. Signal correlation was defined as the average pairwise Pearson’s correlation between trials with the same sound stimulus that do not occur simultaneously (unmatched) from the two cells. Significant signal correlations were identified by comparing the signal correlation of each neuron pair to a distribution computed from an equal number of blank trials 1000 times (two-tailed Z-test, alpha 0.05). Noise correlation was defined as the average pairwise Pearson’s correlation between the residuals of trials with the same sound stimulus that occur simultaneously (matched), computed by subtracting the mean response to that stimulus from each trial for each cell. Significant noise correlations were identified by comparing the noise correlation of each neuron pair to a distribution computed from an equal number of unmatched trials 1000 times (two-tailed Z-test, alpha 0.05). Only significantly correlated pairs were considered for statistical analyses. Pairs with inter-neuronal distance less than 20µm were also not considered to avoid the inclusion of pairs with overlapping ROIs.

### GLM

A generalized linear model (GLM) was fit to neural data using a ridge regression^86^. Calcium traces were randomly divided into testing and training data such that the minimum section of contiguous data contained at least one trial (7 seconds). 20% of data were held out for testing the model fit. The remaining 80% of data were used for training and validation. Of this training/validation set, 80% were used to train and 20% were used to validate. For each validation step, we tested a range of λ parameters (2^0^ - 2^15^) to maximize correlation with testing data. This cross validation was repeated 10x with randomly sectioned testing/validation data to find the λ that produced the maximal data fit. Finally, the model was refit to the entire training and validation dataset, and this refit model was evaluated on the test dataset. Model prediction performance was defined as the correlation coefficient between the predicted and actual calcium responses from the test dataset Regressors for the model included normalized locomotor activity, sound stimuli, and residuals for all other neurons within the field of view, calculated as described above. The final model was the linear combination of these data with (sound and locomotion) or without (residuals of neighboring neurons) delay with respect to event onset.

### Statistical analyses

Linear mixed-effects models^87,88^ were used to analyze data collected through nested experimental designs. For these models, FOV nested within mouse and group were specified as random effects. When data were combined across sound stimuli, neuron or neuron pair (for correlation analyses) nested within FOV, mouse, and group were specified as random effects. Analyses were conducted using MATLAB 2021a and JMP 16 (SAS Institute). The fixed-effect test results are reported with the degrees of freedom denominator approximated for normal data using the Kenward–Roger adjustment. All tests were evaluated at an alpha level of 0.05 and Tukey’s test was used for *post hoc* comparisons unless otherwise noted. Where Bonferroni’s post-hoc correction is used, p-values are reported as the t-test p-value multiplied by the number of comparisons so that the alpha level of 0.05 is held constant. Box and whisker plots represent the median and interquartile range (IQR) from 25-75% (box), and 1.5*IQR (whiskers). Error bars are standard error to the mean (SEM).

## Supplemental information

**Supplement Figure 1:**
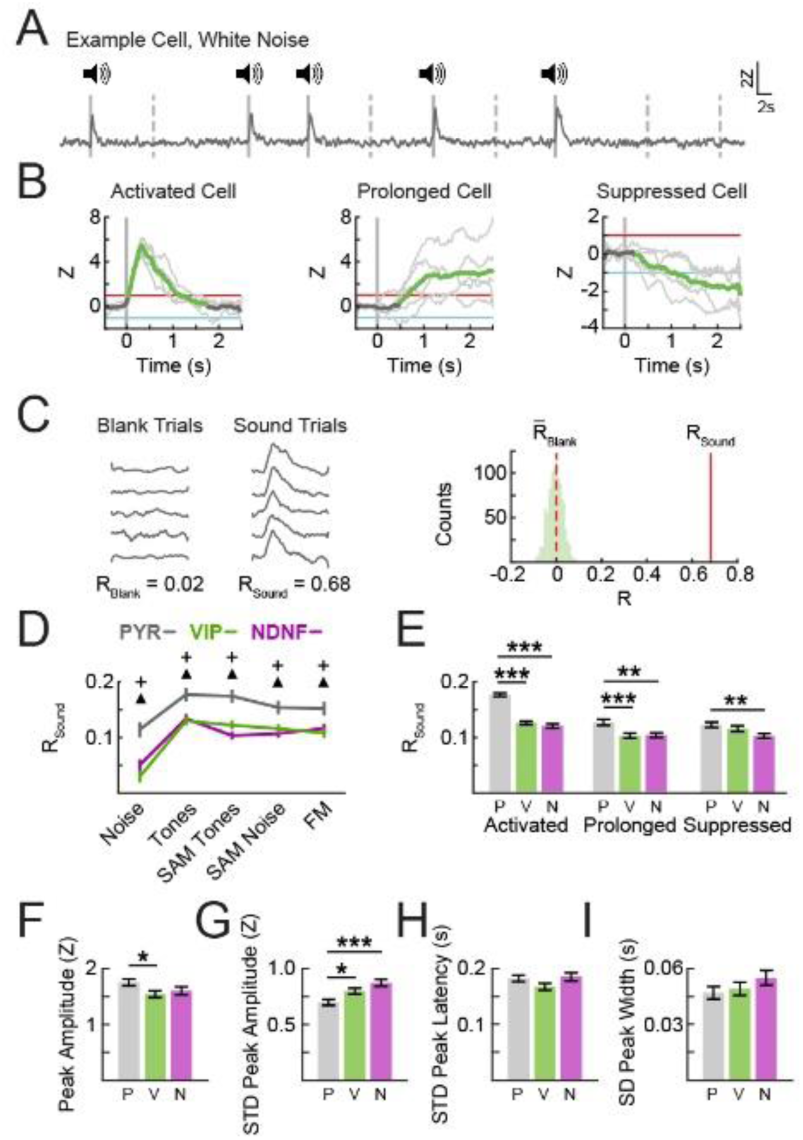
Method for assessing reliable sound responses. **(A)** An example trace of a pyramidal neuron responding to white noise bursts, indicated by solid lines, interleaved with blank (silent) trials, indicated by dotted lines. **(B)** Example *activated*, *prolonged*, and *suppressed* cells. Detectable responses are defined as having an average maximum or minimum response magnitude of at least 1 Z-score (*activated* or *prolonged*) or less than −1 Z-score (*suppressed*). Light gray traces are single trials, dark gray traces are the average response, and green traces are the portion of the average response that does not cross zero (used for calculating response width). **(C)** Example blank trials and sound trials (white noise) from a significantly responsive pyramidal cell. Reliability is calculated as the average pairwise Pearson’s R value between all sound trials (R_Sound_) and compared to a bootstrapped distribution of values computed from blank trials (R_Blank_) where the mean is ^R̅^_Blank_. **(D)** Reliability was greatest for PYR neurons for all stimuli tested: mixed-effects two-way ANOVA, neuron group x stimulus type with post-hoc Bonferroni correction within stimulus type compared to PYR, (F_8,3918_ = 4.0902, p<0.0001), ^+^VIP or ^Δ^NDNF significantly different from PYR, (F_2,75.06_ ≤ 4.2739, p≤0.0175), p<0.0438, n_PYR_ = 694 neurons; n_VIP_ = 773 neurons; n_NDNF_ = 829 neurons. **(E)** PYR neuron reliability was greater than VIP and NDNF interneuron reliability for populations with activated and prolonged response types, and greater than NDNF neurons for populations with suppressed response types: mixed-effects two-way ANOVA, neuron group x response type F_(4,3925)_ = 10.4235, p<0.0001. Post-hoc t-test with Bonferroni correction within response type compared to PYR, (F_2,3776_ ≤ 5.4718, p≤0.0042), ***p<.001, **p<0.0024, n_PYR_ = 694 neurons; n_VIP_ = 773 neurons; n_NDNF_ = 829 neurons. **(F-I)** For *activated* neurons, PYR exhibited the greatest peak response magnitude (F) and lowest standard deviation (SD) in peak magnitude (**G**). The three populations did not differ in the standard deviation of peak width (**H**) or peak latency (**I**). Mixed-effects one-way ANOVAs followed by Tukey’s post-hoc test: peak amplitude (F_2,1404_ = 3.1621, p=0.0426), SD peak amplitude (F_2,1363_ = 9.0687, p=0.0001), SD peak latency (F_2,700.6_ = 2.2624, p=0.1049), SD peak width (F_2,1180_ = 1.2559, p = 0.2852), n_PYR_ = 423 neurons; n_VIP_ = 470 neurons; n_NDNF_ = 374 neurons. ***p<0.001, *p<0.05.

**Supplement Figure 2:**
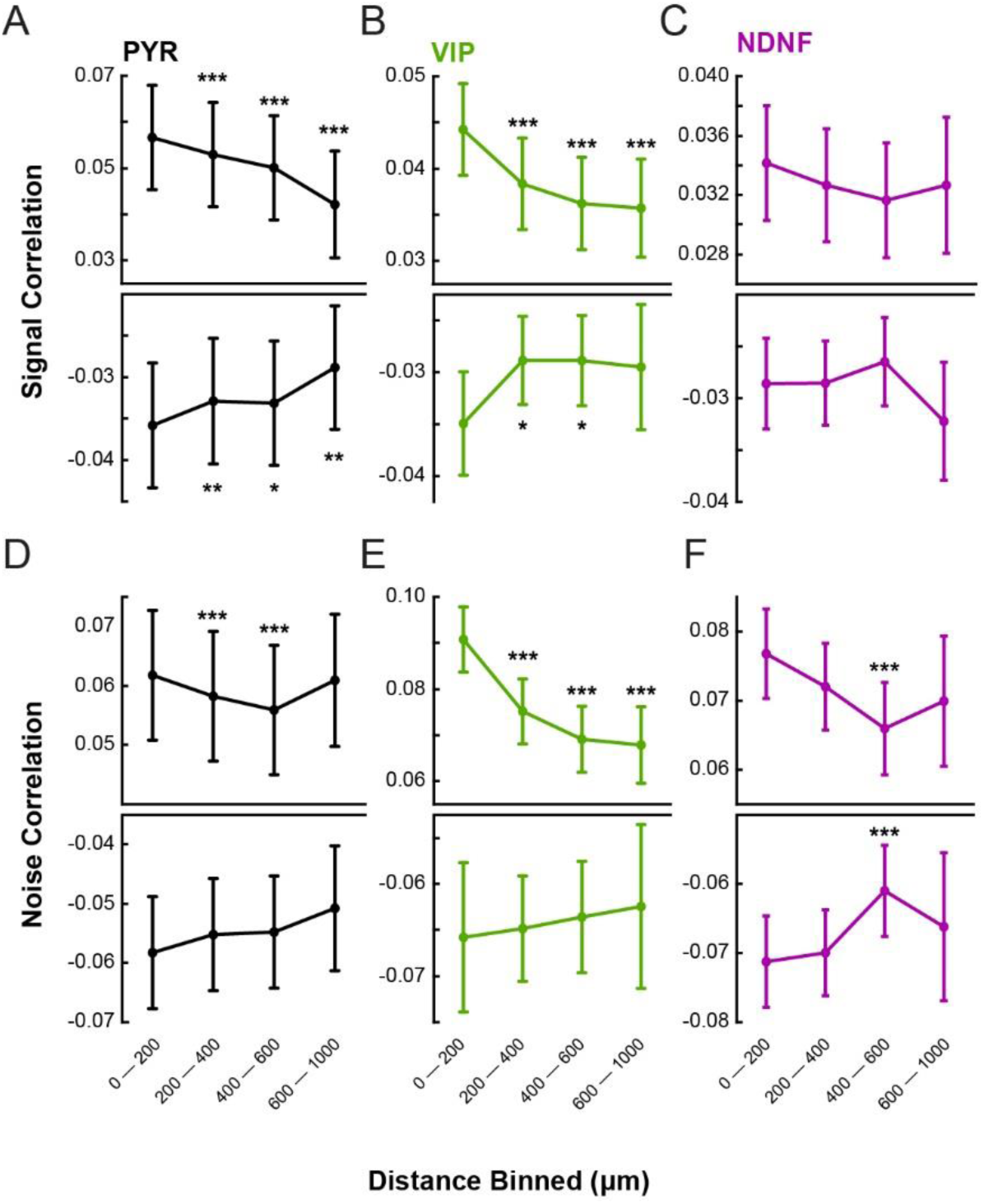
Signal and noise correlations are strongest among neuronal pairs within close spatial proximity. **A-C** Significant pairwise signal correlations for PYR (A), VIP (B), and NDNF (C) neurons binned as a function of distance between neurons within the same field-of-view. **D-F** Significant pairwise noise correlations for PYR (D), VIP (E), and NDNF (F) neurons binned as a function of distance between neurons within the same field-of-view. Mixed-effects one-way ANOVA with Tukey’s post hoc test. ***p<0.001, **p<0.01, *p<0.05. Signal correlations: n_PYR_ = 14861 neuron pairs, n_VIP_ = 5141 neuron pairs, n_NDNF_ = 3268 neuron pairs. Noise correlations: n_PYR_ = 1436 neuron pairs, n_VIP_ = 241 neuron pairs, n_NDNF_ = 944 neuron pairs.

## References

1. Fritz, J.B., Elhilali, M., David, S.V., and Shamma, S.A. (2007). Auditory attention--focusing the searchlight on sound. Curr Opin Neurobiol 17, 437–455. 10.1016/j.conb.2007.07.011.

2. Angeloni, C., and Geffen, M.N. (2018). Contextual modulation of sound processing in the auditory cortex. Curr Opin Neurobiol 49, 8–15. 10.1016/j.conb.2017.10.012.

3. Kuchibhotla, K., and Bathellier, B. (2018). Neural encoding of sensory and behavioral complexity in the auditory cortex. Curr Opin Neurobiol 52, 65–71. 10.1016/j.conb.2018.04.002.

4. Khan, A.G., and Hofer, S.B. (2018). Contextual signals in visual cortex. Curr Opin Neurobiol 52, 131–138. 10.1016/j.conb.2018.05.003.

5. Froemke, R.C., Merzenich, M.M., and Schreiner, C.E. (2007). A synaptic memory trace for cortical receptive field plasticity. Nature 450, 425–429. 10.1038/nature06289.

6. Kuchibhotla, K.V., Gill, J.V., Lindsay, G.W., Papadoyannis, E.S., Field, R.E., Sten, T.A., Miller, K.D., and Froemke, R.C. (2017). Parallel processing by cortical inhibition enables context-dependent behavior. Nature neuroscience 20, 62–71. 10.1038/nn.4436.

7. Blackwell, J.M., and Geffen, M.N. (2017). Progress and challenges for understanding the function of cortical microcircuits in auditory processing. Nature communications 8, 2165. 10.1038/s41467-017-01755-2.

8. Winer, J.A., Miller, L.M., Lee, C.C., and Schreiner, C.E. (2005). Auditory thalamocortical transformation: structure and function. Trends Neurosci 28, 255–263. 10.1016/j.tins.2005.03.009.

9. Lee, S., Kruglikov, I., Huang, Z.J., Fishell, G., and Rudy, B. (2013). A disinhibitory circuit mediates motor integration in the somatosensory cortex. Nature neuroscience 16, 1662–1670. 10.1038/nn.3544.

10. Schroeder, A., Pardi, M.B., Keijser, J., Dalmay, T., Groisman, A.I., Schuman, E.M., Sprekeler, H., and Letzkus, J.J. (2023). Inhibitory top-down projections from zona incerta mediate neocortical memory. Neuron 111, 727–738.e728. 10.1016/j.neuron.2022.12.010.

11. Williams, L.E., and Holtmaat, A. (2019). Higher-Order Thalamocortical Inputs Gate Synaptic Long-Term Potentiation via Disinhibition. Neuron 101, 91–102.e104. 10.1016/j.neuron.2018.10.049.

12. Pardi, M.B., Vogenstahl, J., Dalmay, T., Spanò, T., Pu, D.L., Naumann, L.B., Kretschmer, F., Sprekeler, H., and Letzkus, J.J. (2020). A thalamocortical top-down circuit for associative memory. Science (New York, N.Y.) 370, 844–848. 10.1126/science.abc2399.

13. Cohen-Kashi Malina, K., Tsivourakis, E., Kushinsky, D., Apelblat, D., Shtiglitz, S., Zohar, E., Sokoletsky, M., Tasaka, G.I., Mizrahi, A., Lampl, I., and Spiegel, I. (2021). NDNF interneurons in layer 1 gain-modulate whole cortical columns according to an animal’s behavioral state. Neuron 109, 2150–2164.e2155. 10.1016/j.neuron.2021.05.001.

14. Abs, E., Poorthuis, R.B., Apelblat, D., Muhammad, K., Pardi, M.B., Enke, L., Kushinsky, D., Pu, D.L., Eizinger, M.F., Conzelmann, K.K., et al. (2018). Learning-Related Plasticity in Dendrite-Targeting Layer 1 Interneurons. Neuron 100, 684–699.e686. 10.1016/j.neuron.2018.09.001.

15. Takesian, A.E., Bogart, L.J., Lichtman, J.W., and Hensch, T.K. (2018). Inhibitory circuit gating of auditory critical-period plasticity. Nature neuroscience 21, 218–227. 10.1038/s41593-017-0064-2.

16. Hartung, J., and Letzkus, J.J. (2021). Inhibitory plasticity in layer 1 - dynamic gatekeeper of neocortical associations. Curr Opin Neurobiol 67, 26–33. 10.1016/j.conb.2020.06.003.

17. Huang, S., Wu, S.J., Sansone, G., Ibrahim, L.A., and Fishell, G. (2024). Layer 1 neocortex: Gating and integrating multidimensional signals. Neuron 112, 184–200. 10.1016/j.neuron.2023.09.041.

18. Cruz-Martín, A., El-Danaf, R.N., Osakada, F., Sriram, B., Dhande, O.S., Nguyen, P.L., Callaway, E.M., Ghosh, A., and Huberman, A.D. (2014). A dedicated circuit links direction-selective retinal ganglion cells to the primary visual cortex. Nature 507, 358–361. 10.1038/nature12989.

19. Roth, M.M., Dahmen, J.C., Muir, D.R., Imhof, F., Martini, F.J., and Hofer, S.B. (2016). Thalamic nuclei convey diverse contextual information to layer 1 of visual cortex. Nature neuroscience 19, 299–307. 10.1038/nn.4197.

20. Zhu, F., Elnozahy, S., Lawlor, J., and Kuchibhotla, K.V. (2023). The cholinergic basal forebrain provides a parallel channel for state-dependent sensory signaling to auditory cortex. Nature neuroscience 26, 810–819. 10.1038/s41593-023-01289-5.

21. Letzkus, J.J., Wolff, S.B., Meyer, E.M., Tovote, P., Courtin, J., Herry, C., and Luthi, A. (2011). A disinhibitory microcircuit for associative fear learning in the auditory cortex. Nature 480, 331–335. 10.1038/nature10674.

22. Trottier, S., Evrard, B., Vignal, J.P., Scarabin, J.M., and Chauvel, P. (1996). The serotonergic innervation of the cerebral cortex in man and its changes in focal cortical dysplasia. Epilepsy Res 25, 79–106. 10.1016/0920-1211(96)00033-2.

23. Kajstura, T.J., Dougherty, S.E., and Linden, D.J. (2018). Serotonin axons in the neocortex of the adult female mouse regrow after traumatic brain injury. J Neurosci Res 96, 512–526. 10.1002/jnr.24059.

24. Mechawar, N., Cozzari, C., and Descarries, L. (2000). Cholinergic innervation in adult rat cerebral cortex: a quantitative immunocytochemical description. The Journal of comparative neurology 428, 305–318. 10.1002/1096-9861(20001211)428:2<305::aid-cne9>3.0.co;2-y.

25. Eggermann, E., Kremer, Y., Crochet, S., and Petersen, C.C.H. (2014). Cholinergic signals in mouse barrel cortex during active whisker sensing. Cell reports 9, 1654–1660. 10.1016/j.celrep.2014.11.005.

26. Mowery, T.M., Kotak, V.C., and Sanes, D.H. (2016). The onset of visual experience gates auditory cortex critical periods. Nature communications 7, 10416. 10.1038/ncomms10416.

27. Sun, W., Tang, P., Liang, Y., Li, J., Feng, J., Zhang, N., Lu, D., He, J., and Chen, X. (2022). The anterior cingulate cortex directly enhances auditory cortical responses in air-puffing-facilitated flight behavior. Cell reports 38, 110506. 10.1016/j.celrep.2022.110506.

28. Ibrahim, L.A., Mesik, L., Ji, X.Y., Fang, Q., Li, H.F., Li, Y.T., Zingg, B., Zhang, L.I., and Tao, H.W. (2016). Cross-Modality Sharpening of Visual Cortical Processing through Layer-1-Mediated Inhibition and Disinhibition. Neuron 89, 1031–1045. 10.1016/j.neuron.2016.01.027.

29. Leinweber, M., Ward, D.R., Sobczak, J.M., Attinger, A., and Keller, G.B. (2017). A Sensorimotor Circuit in Mouse Cortex for Visual Flow Predictions. Neuron 95, 1420–1432.e1425. 10.1016/j.neuron.2017.08.036.

30. Ledderose, J.M.T., Zolnik, T.A., Toumazou, M., Trimbuch, T., Rosenmund, C., Eickholt, B.J., Jaeger, D., Larkum, M.E., and Sachdev, R.N.S. (2023). Layer 1 of somatosensory cortex: an important site for input to a tiny cortical compartment. Cerebral cortex (New York, N.Y. : 1991) 33, 11354–11372. 10.1093/cercor/bhad371.

31. Yang, Y., Liu, D.Q., Huang, W., Deng, J., Sun, Y., Zuo, Y., and Poo, M.M. (2018). Author Correction: Selective synaptic remodeling of amygdalocortical connections associated with fear memory. Nature neuroscience 21, 1137. 10.1038/s41593-018-0180-7.

32. Wall, N.R., De La Parra, M., Sorokin, J.M., Taniguchi, H., Huang, Z.J., and Callaway, E.M. (2016). Brain-Wide Maps of Synaptic Input to Cortical Interneurons. J Neurosci 36, 4000–4009. 10.1523/jneurosci.3967-15.2016.

33. Vattino, L.G., MacGregor, C.P., Liu, C.J., Sweeney, C.G., and Takesian, A.E. (2024). Primary auditory thalamus relays directly to cortical layer 1 interneurons. bioRxiv, 2024.2007.2016.603741. 10.1101/2024.07.16.603741.

34. Schuman, B., Machold, R.P., Hashikawa, Y., Fuzik, J., Fishell, G.J., and Rudy, B. (2019). Four Unique Interneuron Populations Reside in Neocortical Layer 1. J Neurosci 39, 125–139. 10.1523/jneurosci.1613-18.2018.

35. Fu, Y., Kaneko, M., Tang, Y., Alvarez-Buylla, A., and Stryker, M.P. (2015). A cortical disinhibitory circuit for enhancing adult plasticity. eLife 4, e05558. 10.7554/eLife.05558.

36. Fu, Y., Tucciarone, J.M., Espinosa, J.S., Sheng, N., Darcy, D.P., Nicoll, R.A., Huang, Z.J., and Stryker, M.P. (2014). A cortical circuit for gain control by behavioral state. Cell 156, 1139–1152. 10.1016/j.cell.2014.01.050.

37. Pi, H.J., Hangya, B., Kvitsiani, D., Sanders, J.I., Huang, Z.J., and Kepecs, A. (2013). Cortical interneurons that specialize in disinhibitory control. Nature 503, 521–524. 10.1038/nature12676.

38. Melzer, S., Newmark, E.R., Mizuno, G.O., Hyun, M., Philson, A.C., Quiroli, E., Righetti, B., Gregory, M.R., Huang, K.W., Levasseur, J., et al. (2021). Bombesin-like peptide recruits disinhibitory cortical circuits and enhances fear memories. Cell 184, 5622–5634.e5625. 10.1016/j.cell.2021.09.013.

39. Szadai, Z., Pi, H.J., Chevy, Q., Ócsai, K., Albeanu, D.F., Chiovini, B., Szalay, G., Katona, G., Kepecs, A., and Rózsa, B. (2022). Cortex-wide response mode of VIP-expressing inhibitory neurons by reward and punishment. eLife 11. 10.7554/eLife.78815.

40. Smith, P.H., Uhlrich, D.J., Manning, K.A., and Banks, M.I. (2012). Thalamocortical projections to rat auditory cortex from the ventral and dorsal divisions of the medial geniculate nucleus. The Journal of comparative neurology 520, 34–51. 10.1002/cne.22682.

41. Ji, X.Y., Zingg, B., Mesik, L., Xiao, Z., Zhang, L.I., and Tao, H.W. (2016). Thalamocortical Innervation Pattern in Mouse Auditory and Visual Cortex: Laminar and Cell-Type Specificity. Cerebral cortex (New York, N.Y. : 1991) 26, 2612–2625. 10.1093/cercor/bhv099.

42. Hackett, T.A., Clause, A.R., Takahata, T., Hackett, N.J., and Polley, D.B. (2016). Differential maturation of vesicular glutamate and GABA transporter expression in the mouse auditory forebrain during the first weeks of hearing. Brain Struct Funct 221, 2619–2673. 10.1007/s00429-015-1062-3.

43. Vasquez-Lopez, S.A., Weissenberger, Y., Lohse, M., Keating, P., King, A.J., and Dahmen, J.C. (2017). Thalamic input to auditory cortex is locally heterogeneous but globally tonotopic. eLife 6. 10.7554/eLife.25141.

44. Poorthuis, R.B., Muhammad, K., Wang, M., Verhoog, M.B., Junek, S., Wrana, A., Mansvelder, H.D., and Letzkus, J.J. (2018). Rapid Neuromodulation of Layer 1 Interneurons in Human Neocortex. Cell reports 23, 951–958. 10.1016/j.celrep.2018.03.111.

45. Mesik, L., Ma, W.P., Li, L.Y., Ibrahim, L.A., Huang, Z.J., Zhang, L.I., and Tao, H.W. (2015). Functional response properties of VIP-expressing inhibitory neurons in mouse visual and auditory cortex. Front Neural Circuits 9, 22. 10.3389/fncir.2015.00022.

46. Tischbirek, C.H., Noda, T., Tohmi, M., Birkner, A., Nelken, I., and Konnerth, A. (2019). In Vivo Functional Mapping of a Cortical Column at Single-Neuron Resolution. Cell reports 27, 1319–1326.e1315. 10.1016/j.celrep.2019.04.007.

47. Romero, S., Hight, A.E., Clayton, K.K., Resnik, J., Williamson, R.S., Hancock, K.E., and Polley, D.B. (2020). Cellular and Widefield Imaging of Sound Frequency Organization in Primary and Higher Order Fields of the Mouse Auditory Cortex. Cerebral cortex (New York, N.Y. : 1991) 30, 1603–1622. 10.1093/cercor/bhz190.

48. Winkowski, D.E., and Kanold, P.O. (2013). Laminar transformation of frequency organization in auditory cortex. J Neurosci 33, 1498–1508. 10.1523/jneurosci.3101-12.2013.

49. Issa, J.B., Haeffele, B.D., Agarwal, A., Bergles, D.E., Young, E.D., and Yue, D.T. (2014). Multiscale optical Ca2+ imaging of tonal organization in mouse auditory cortex. Neuron 83, 944–959. 10.1016/j.neuron.2014.07.009.

50. Panniello, M., King, A.J., Dahmen, J.C., and Walker, K.M.M. (2018). Local and Global Spatial Organization of Interaural Level Difference and Frequency Preferences in Auditory Cortex. Cerebral cortex (New York, N.Y. : 1991) 28, 350–369. 10.1093/cercor/bhx295.

51. Liu, J., Whiteway, M.R., Sheikhattar, A., Butts, D.A., Babadi, B., and Kanold, P.O. (2019). Parallel Processing of Sound Dynamics across Mouse Auditory Cortex via Spatially Patterned Thalamic Inputs and Distinct Areal Intracortical Circuits. Cell reports 27, 872–885.e877. 10.1016/j.celrep.2019.03.069.

52. Guo, W., Chambers, A.R., Darrow, K.N., Hancock, K.E., Shinn-Cunningham, B.G., and Polley, D.B. (2012). Robustness of cortical topography across fields, laminae, anesthetic states, and neurophysiological signal types. J Neurosci 32, 9159–9172. 10.1523/jneurosci.0065-12.2012.

53. Schneider, D.M., Nelson, A., and Mooney, R. (2014). A synaptic and circuit basis for corollary discharge in the auditory cortex. Nature 513, 189–194. 10.1038/nature13724.

54. McGinley, M.J., David, S.V., and McCormick, D.A. (2015). Cortical Membrane Potential Signature of Optimal States for Sensory Signal Detection. Neuron 87, 179–192. 10.1016/j.neuron.2015.05.038.

55. Zhou, M., Liang, F., Xiong, X.R., Li, L., Li, H., Xiao, Z., Tao, H.W., and Zhang, L.I. (2014). Scaling down of balanced excitation and inhibition by active behavioral states in auditory cortex. Nature neuroscience 17, 841–850. 10.1038/nn.3701.

56. Faisal, A.A., Selen, L.P., and Wolpert, D.M. (2008). Noise in the nervous system. Nature reviews. Neuroscience 9, 292–303. 10.1038/nrn2258.

57. Doiron, B., Litwin-Kumar, A., Rosenbaum, R., Ocker, G.K., and Josić, K. (2016). The mechanics of state-dependent neural correlations. Nature neuroscience 19, 383–393. 10.1038/nn.4242.

58. Shi, Y.L., Steinmetz, N.A., Moore, T., Boahen, K., and Engel, T.A. (2022). Cortical state dynamics and selective attention define the spatial pattern of correlated variability in neocortex. Nature communications 13, 44. 10.1038/s41467-021-27724-4.

59. Ko, H., Hofer, S.B., Pichler, B., Buchanan, K.A., Sjöström, P.J., and Mrsic-Flogel, T.D. (2011). Functional specificity of local synaptic connections in neocortical networks. Nature 473, 87–91. 10.1038/nature09880.

60. Yao, X.H., Wang, M., He, X.N., He, F., Zhang, S.Q., Lu, W., Qiu, Z.L., and Yu, Y.C. (2016). Electrical coupling regulates layer 1 interneuron microcircuit formation in the neocortex. Nature communications 7, 12229. 10.1038/ncomms12229.

61. Fan, L.Z., Kheifets, S., Böhm, U.L., Wu, H., Piatkevich, K.D., Xie, M.E., Parot, V., Ha, Y., Evans, K.E., Boyden, E.S., et al. (2020). All-Optical Electrophysiology Reveals the Role of Lateral Inhibition in Sensory Processing in Cortical Layer 1. Cell 180, 521–535.e518. 10.1016/j.cell.2020.01.001.

62. Grimsley, J.M., Monaghan, J.J., and Wenstrup, J.J. (2011). Development of social vocalizations in mice. PLoS One 6, e17460. 10.1371/journal.pone.0017460.

63. Holy, T.E., and Guo, Z. (2005). Ultrasonic songs of male mice. PLoS Biol 3, e386. 10.1371/journal.pbio.0030386.

64. Portfors, C.V. (2007). Types and functions of ultrasonic vocalizations in laboratory rats and mice. J Am Assoc Lab Anim Sci 46, 28–34.

65. Liu, R.C., Miller, K.D., Merzenich, M.M., and Schreiner, C.E. (2003). Acoustic variability and distinguishability among mouse ultrasound vocalizations. J Acoust Soc Am 114, 3412–3422. 10.1121/1.1623787.

66. Hartung, J., Schroeder, A., Péréz Vázquez, R.A., Poorthuis, R.B., and Letzkus, J.J. (2024). Layer 1 NDNF interneurons are specialized top-down master regulators of cortical circuits. Cell reports 43, 114212. 10.1016/j.celrep.2024.114212.

67. Kerlin, A.M., Andermann, M.L., Berezovskii, V.K., and Reid, R.C. (2010). Broadly tuned response properties of diverse inhibitory neuron subtypes in mouse visual cortex. Neuron 67, 858–871. 10.1016/j.neuron.2010.08.002.

68. Ebrahimi, S., Lecoq, J., Rumyantsev, O., Tasci, T., Zhang, Y., Irimia, C., Li, J., Ganguli, S., and Schnitzer, M.J. (2022). Emergent reliability in sensory cortical coding and inter-area communication. Nature 605, 713–721. 10.1038/s41586-022-04724-y.

69. Zhu, Y., Qiao, W., Liu, K., Zhong, H., and Yao, H. (2015). Control of response reliability by parvalbumin-expressing interneurons in visual cortex. Nature communications 6, 6802. 10.1038/ncomms7802.

70. Rikhye, R.V., Yildirim, M., Hu, M., Breton-Provencher, V., and Sur, M. (2021). Reliable Sensory Processing in Mouse Visual Cortex through Cooperative Interactions between Somatostatin and Parvalbumin Interneurons. J Neurosci 41, 8761–8778. 10.1523/jneurosci.3176-20.2021.

71. Kara, P., Reinagel, P., and Reid, R.C. (2000). Low response variability in simultaneously recorded retinal, thalamic, and cortical neurons. Neuron 27, 635–646. 10.1016/s0896-6273(00)00072-6.

72. Willmore, B.D., Mazer, J.A., and Gallant, J.L. (2011). Sparse coding in striate and extrastriate visual cortex. J Neurophysiol 105, 2907–2919. 10.1152/jn.00594.2010.

73. Olshausen, B.A., and Field, D.J. (2004). Sparse coding of sensory inputs. Curr Opin Neurobiol 14, 481–487. 10.1016/j.conb.2004.07.007.

74. Zhang, C., Zhang, D., and Stepanyants, A. (2021). Noise in Neurons and Synapses Enables Reliable Associative Memory Storage in Local Cortical Circuits. eNeuro 8. 10.1523/eneuro.0302-20.2020.

75. Benedetti, B.L., Glazewski, S., and Barth, A.L. (2009). Reliable and precise neuronal firing during sensory plasticity in superficial layers of primary somatosensory cortex. J Neurosci 29, 11817–11827. 10.1523/jneurosci.3431-09.2009.

76. Caruso, V.C., Mohl, J.T., Glynn, C., Lee, J., Willett, S.M., Zaman, A., Ebihara, A.F., Estrada, R., Freiwald, W.A., Tokdar, S.T., and Groh, J.M. (2018). Single neurons may encode simultaneous stimuli by switching between activity patterns. Nature communications 9, 2715. 10.1038/s41467-018-05121-8.

77. Huang, S., Rizzo, D., Wu, S.J., Xu, Q., Ziane, L., Alghamdi, N., Stafford, D.A., Daigle, T.L., Tasic, B., Zeng, H., et al. (2024). Neurogliaform Cells Exhibit Laminar-specific Responses in the Visual Cortex and Modulate Behavioral State-dependent Cortical Activity. bioRxiv, 2024.2006.2005.597539. 10.1101/2024.06.05.597539.

78. Granger, A.J., Wang, W., Robertson, K., El-Rifai, M., Zanello, A.F., Bistrong, K., Saunders, A., Chow, B.W., Nuñez, V., Turrero García, M., et al. (2020). Cortical ChAT(+) neurons co-transmit acetylcholine and GABA in a target- and brain-region-specific manner. eLife 9. 10.7554/eLife.57749.

79. Obermayer, J., Luchicchi, A., Heistek, T.S., de Kloet, S.F., Terra, H., Bruinsma, B., Mnie-Filali, O., Kortleven, C., Galakhova, A.A., Khalil, A.J., et al. (2020). Author Correction: Prefrontal cortical ChAT-VIP interneurons provide local excitation by cholinergic synaptic transmission and control attention. Nature communications 11, 794. 10.1038/s41467-020-14315-y.

80. Dudai, A., Yayon, N., Lerner, V., Tasaka, G.I., Deitcher, Y., Gorfine, K., Niederhoffer, N., Mizrahi, A., Soreq, H., and London, M. (2020). Barrel cortex VIP/ChAT interneurons suppress sensory responses in vivo. PLoS Biol 18, e3000613. 10.1371/journal.pbio.3000613.

81. Jiang, X., Wang, G., Lee, A.J., Stornetta, R.L., and Zhu, J.J. (2013). The organization of two new cortical interneuronal circuits. Nature neuroscience 16, 210–218. 10.1038/nn.3305.

82. Karnani, M.M., Jackson, J., Ayzenshtat, I., Hamzehei Sichani, A., Manoocheri, K., Kim, S., and Yuste, R. (2016). Opening Holes in the Blanket of Inhibition: Localized Lateral Disinhibition by VIP Interneurons. J Neurosci 36, 3471–3480. 10.1523/jneurosci.3646-15.2016.

83. Jackson, J., Ayzenshtat, I., Karnani, M.M., and Yuste, R. (2016). VIP+ interneurons control neocortical activity across brain states. J Neurophysiol 115, 3008–3017. 10.1152/jn.01124.2015.

84. Cruikshank, S.J., Rose, H.J., and Metherate, R. (2002). Auditory thalamocortical synaptic transmission in vitro. J Neurophysiol 87, 361–384. 10.1152/jn.00549.2001.

85. Pachitariu, M., Stringer, C., Dipoppa, M., Schröder, S., Rossi, L.F., Dalgleish, H., Carandini, M., and Harris, K.D. (2017). Suite2p: beyond 10,000 neurons with standard two-photon microscopy. bioRxiv, 061507. 10.1101/061507.

86. Pinto, L., and Dan, Y. (2015). Cell-Type-Specific Activity in Prefrontal Cortex during Goal-Directed Behavior. Neuron 87, 437–450. 10.1016/j.neuron.2015.06.021.

87. Reed, J.L., and Kaas, J.H. (2010). Statistical analysis of large-scale neuronal recording data. Neural Netw 23, 673–684. 10.1016/j.neunet.2010.04.005.

88. Aarts, E., Verhage, M., Veenvliet, J.V., Dolan, C.V., and van der Sluis, S. (2014). A solution to dependency: using multilevel analysis to accommodate nested data. Nature neuroscience 17, 491–496. 10.1038/nn.3648.

